# Scene-selectivity in CA1/subicular complex: Multivoxel pattern analysis at 7T

**DOI:** 10.1101/2023.10.02.560435

**Authors:** Marie-Lucie Read, Samuel C. Berry, Kim S. Graham, Natalie L. Voets, Jiaxiang Zhang, John P. Aggleton, Andrew D. Lawrence, Carl J. Hodgetts

**Author notes:** Joint first authors (Marie-Lucie Read, Samuel C. Berry).

## Abstract

Prior univariate functional magnetic resonance imaging (fMRI) studies in humans suggest that the anteromedial subicular complex of the hippocampus is a hub for scene-based cognition. However, it is possible that univariate approaches were not sufficiently sensitive to detect scene-related activity in other subfields that have been implicated in spatial processing (e.g., CA1). Further, as connectivity-based functional gradients in the hippocampus do not respect classical subfield boundary definitions, category sensitivity may be distributed across anatomical subfields. Region-of-interest approaches, therefore, may limit our ability to observe category selectivity across discrete subfield boundaries. To address these issues, we applied searchlight multivariate pattern analysis to 7T fMRI data of healthy adults who undertook a simultaneous visual odd-one-out discrimination task for scene and non-scene (including face) visual stimuli, hypothesising that scene classification would be possible in multiple hippocampal regions within, but not constrained to, anteromedial subicular complex and CA1. Indeed, we found that the scene-selective searchlight map overlapped not only with anteromedial subicular complex (distal subiculum, pre/para subiculum), but also inferior CA1, alongside retrosplenial and parahippocampal cortices. Probabilistic overlap maps revealed gradients of scene category selectivity, with the strongest overlap located in the medial hippocampus, converging with searchlight findings. This was contrasted with gradients of face category selectivity, which had stronger overlap in more lateral hippocampus, supporting ideas of parallel processing streams for these two categories. Our work helps to map the scene, in contrast to, face processing networks within, and connected to, the human hippocampus.

## 1. Introduction

There is increasing evidence that the human hippocampus has roles in behaviours beyond purely-mnemonic cognitive functions, including complex visual perception and imagination (Aly et al., 2013; Graham et al., 2010; Hodgetts et al., 2015; Lee et al., 2013; Martin & Barense, 2023). While the precise role played by the hippocampus in supporting such functions is debated (Mayes, Montaldi & Migo, 2007; Turk-Browne, 2019), there is increasing support for the idea that scenes are central to hippocampal information processing (Maguire & Mullally, 2013; Murray et al., 2017, 2018; Zeidman, Mullally, et al., 2015). Indeed, scene-selective impairments following hippocampal damage are not only seen on memory tasks (Bird et al., 2008; Hartley et al., 2007; Taylor et al., 2007;) but also on tasks of complex visual perception (Graham et al., 2010; Lee et al., 2012; Erez et al., 2013; Martin & Barense, 2023). Some of the strongest evidence for this comes from ‘oddity’ simultaneous visual discrimination paradigms (Lee et al., 2006; Lee et al., 2005), modified from the nonhuman animal literature (e.g., see Buckley et al., 2001). In these tasks, participants are presented with an array of scene or non-scene stimuli on each trial (typically 3 or 4 items per trial) and are required to select the odd-one-out as quickly and as accurately as possible (see Figure 1). Critically, this task has almost no mnemonic demand, as stimuli are never repeated during the task, and all compared items are presented concurrently. Using a four-choice version of this task, Lee and colleagues (Lee et al., 2006; Lee et al., 2005) found that individuals with hippocampal lesions were impaired when having to identify incongruent scenes but not other classes of stimuli (e.g., faces, objects and colour). This scene-selective impairment was observed when scenes were presented from different viewpoints, but not when the scenes were shown from the same viewpoint. In contrast, patients with both hippocampal and perirhinal cortex (PRC) damage demonstrated significant deficits for both scenes and faces shown from different views but performed normally for scenes and faces shown from the same view (Lee et al., 2005; see also Gardette et al., 2023).

**Figure 1.**
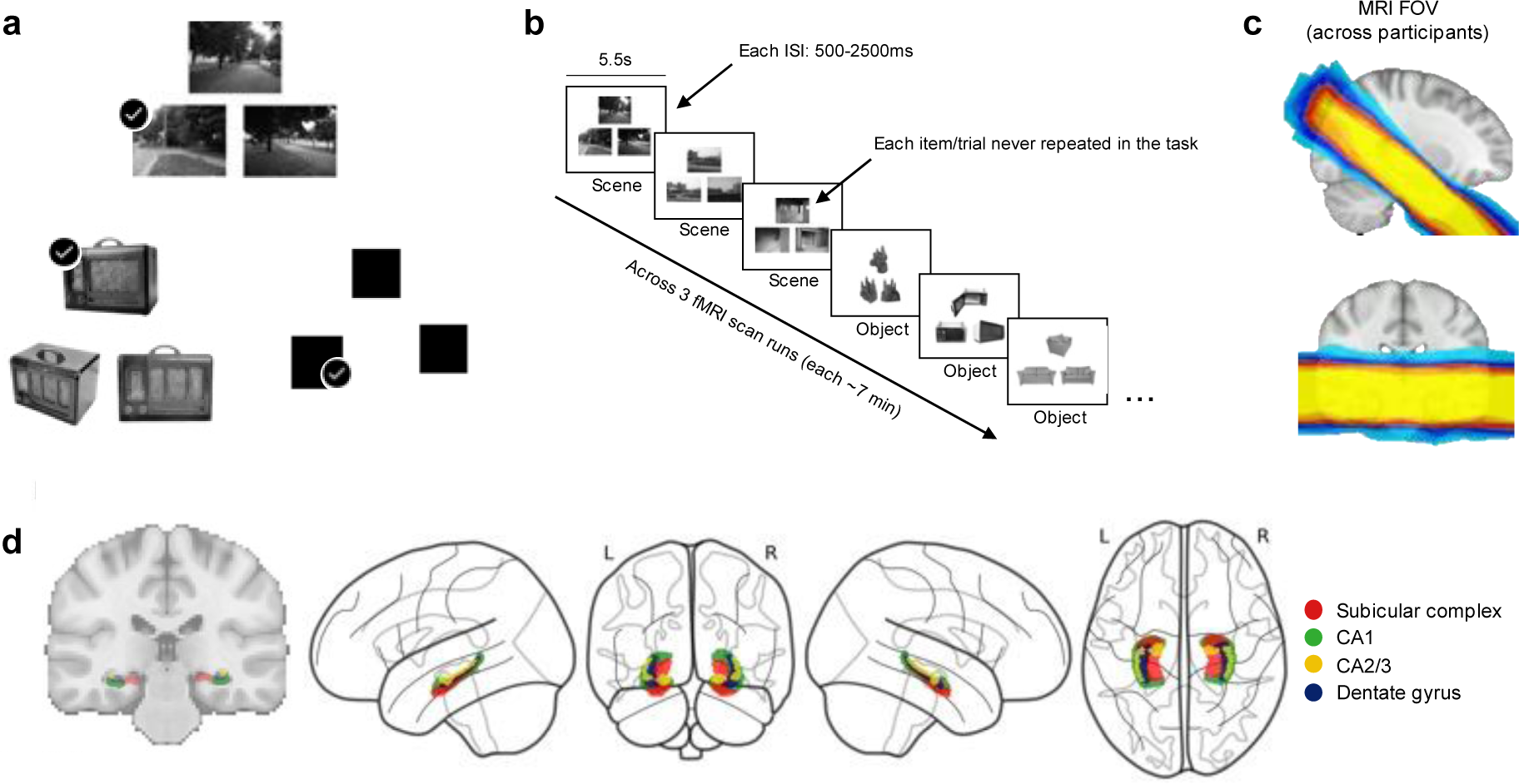
The oddity behavioral task, the partial FOV, and probabilistic hippocampal subfield ROIs in 1mm MNI template space. **a)** Examples of object, scene, and size trials, from top left to bottom right (face trials are not shown due to confidentiality rules, though see Hodgetts et al., 2017 for an example). All stimuli were trial unique. **b)** An illustration of the oddity task procedure. Three trials of the same category were shown sequentially, creating mini-blocks (e.g., here three scene and three face trials are shown). Each trial was presented for 5.5s and trials were separated by a jittered inter-stimulus interval (ISI) of 0.5-2.5s. **d)** The partial FOV of each participant overlaid in MNI template space. Warmer colors indicate higher overlap between participants. The central yellow area indicates where all participant FOVs. overlapped (rendered using FSLeyes; McCarthy (2018). **d)** The left image shows a coronal slice through the hippocampal subfields on a 1mm MNI standard brain. The right images show the same ROIs on a glass brain (rendered using Nilearn for Python; see (Abraham et al., 2014). The ROIs are colored according to the key on the right. L: Left, R: Right.

Functional magnetic resonance imaging (fMRI) studies in healthy participants have attempted to further elucidate the nature and topography of hippocampal scene processing. Earlier studies, for example, primarily reported involvement of the posterior hippocampus in scene perception, including during oddity discrimination tasks (Barense et al., 2010; Lee et al., 2008). However, more recent studies have since suggested that an anteromedial hippocampal region may be a key ‘hub’ for scene-based cognition, including ‘compositional’ aspects of scene perception that require construction of a global scene ‘model’, such as matching scenes from different viewpoints (Baldassano et al., 2016; Hodgetts et al., 2016; Zeidman, Lutti, et al., 2015; Zeidman & Maguire, 2016; Zeidman, Mullally, et al., 2015). Importantly, this has been shown in larger samples and at different levels of spatial smoothing, addressing prior concerns that posterior hippocampal activations reflected signal bleed from adjacent category-selective regions of posterior parahippocampal cortex (Hodgetts et al., 2016). Such variation in scene activation foci across tasks (and indeed individuals) within the hippocampus is unsurprising given that it is not a single homogeneous structure. The hippocampus can be organised into subfields (e.g., CA1, CA2, CA3, dentate gyrus and subiculum) based on cytoarchitectonic features (Schultz & Engelhardt, 2014), and further functional subdivisions may be proposed along both the anterior-posterior and medial-lateral axes based on connectivity patterns (Aggleton & Christiansen, 2015; Christiansen et al., 2017; Dalton et al., 2019; Ezama et al., 2021; Fritch et al., 2021), microstructure (Genon et al., 2021), gene expression patterns (Vogel et al., 2020), and cellular dynamics (e.g., the receptive field size of place cells) (Brunec et al., 2018; Strange et al., 2014). It should be noted that human hippocampal anatomical terms ‘medial’ and ‘lateral’ mostly align with animal hippocampal terms ‘distal’ and ‘proximal’ (see footnote^1^).

Given the methodological challenges associated with characterising such fine-grained functional and/or structural variation within the hippocampus using conventional 3T MRI, further studies have adopted ultra-high field MRI methods to study subfield contributions to scene-related cognition. For example, we previously used high-resolution fMRI (1.2 mm isotropic voxels) to investigate the contribution of different hippocampal subfields (CA1, CA2/3, dentate gyrus, and the subiculum) to scene perception using a 3-choice oddity paradigm (Hodgetts et al., 2017). We found that the subiculum was the only hippocampal substructure to show an increased univariate BOLD response during the perceptual discrimination of scenes, but not face or objects. Further analysis showed that this was solely evident in the anteromedial aspect of the subiculum, a region that we refer to as the anteromedial subicular complex henceforth^2^. This finding thus refines the results of previous 3T fMRI studies (Hodgetts et al., 2016; Lee et al., 2013; Zeidman, Mullally, et al., 2015), by suggesting that the anteromedial hippocampus, and specifically the anteromedial *subicular complex*, may play a unique role in complex scene perception.

While this study and others (e.g. McCormick et al., 2021a; Gardette et al., 2022) have provided important insights into subregional functional variation in the hippocampus, as well as the potentially unique importance of the anteromedial hippocampus/subicular complex in scene processing (Zeidman & Maguire, 2016; Zeidman, Mullally, et al., 2015), these approaches are limited in several respects. First, as noted above, functional gradients in the hippocampus (whether arising via patterns of extrinsic connectivity, gene expression, or cellular dynamics) have been shown to not respect classical subfield boundary definitions (Aggleton & Christiansen, 2015; Dalton et al., 2022; Dalton et al., 2019). Electrophysiological studies in animals, for instance, suggest that spatial information in CA1 is not represented uniformly along its transverse axis, but more strongly represented in proximal CA1 (Henriksen et al., 2010; Ng et al., 2018). Notably, this proximal CA1 region (defined as the CA1 region which borders CA2; Amaral et al., 1991) is also preferentially connected to distal subiculum (defined in the animal literature as the subiculum proper region which borders the pre-subiculum) – a region which likely overlaps with the anteromedial scene region identified in human fMRI studies (Hodgetts et al., 2016; Lee et al., 2013; Zeidman, Mullally, et al., 2015). This spatially modulated circuit is also seemingly preserved through both entorhinal and parahippocampal cortices (Aggleton, 2012; Aggleton & Christiansen, 2015; Gigg, 2006; Witter & Amaral, 2021). Extending this work to human data, a recent study from Dalton et al. (2022) applied tract density mapping to diffusion MRI data, and found that areas of high tract endpoint density tended to extend across classical subfield boundaries (e.g., across the distal subiculum-proximal presubiculum border, and across proximal subiculum and distal CA1 border). This finding also resonates with functional connectivity studies that show long- and transverse-axis gradients of scene selectivity, not only within entorhinal cortex, but also subicular complex (Grande et al., 2022; Maass et al., 2015; Navarro Schröder et al., 2015; see also Schultz, Sommer & Peters, 2015). Overall, therefore, scene selectivity is likely to be distributed across, and vary within, the borders of cytoarchitecturally defined subfields – a pattern that anatomically-defined subfield ROI-based analyses cannot capture without drawing arbitrary sub-divisions (e.g., splitting subicular complex into medial and lateral components in Hodgetts et al., 2017). Further work is required, therefore, to examine scene-selectivity using methods that enable informational content to be mapped within - and across - ROI boundaries.

Second, most previous studies examining hippocampal scene processing (including at high-resolution) have relied on standard univariate analysis approaches that consider only differences in mean activation magnitude as a marker of category selectivity (e.g., Hodgetts et al., 2015; Hodgetts et al., 2016; Hodgetts et al., 2017; Zeidman, Lutti, et al., 2015; Zeidman, Mullally, et al., 2015; but see Liang et al., 2013, Bainbridge, Hall & Baker, 2020). Critically, it is possible that that nonmaximal response patterns carry important (and reliable) category-selective information that cannot be detected using standard methods (Haxby, 2012; Haxby et al., 2014). Spatial/scene information within hippocampal subregions is also likely to be encoded within a distributed neural population code that cannot be captured reliably at the level of individual neurons (Leutgeb et al., 2007; Stefanini et al., 2020) – a principle that may well hold at the level of fMRI voxels (Guest & Love, 2017; Kriegeskorte et al., 2008). Aside from this, there is also considerable evidence (particularly from nonhuman species) that hippocampal subregions outside the subiculum support aspects of spatial and/or scene processing (e.g., see Leutgeb et al., 2007, O’Keefe & Recce, 1993 and Oliva et al., 2016 on neurones displaying location-specific firing fields and theta phase precession in CA1, CA2, CA3 and the DG; and Robertson et al., 1998 and Rolls, 1999, 2023 on spatial view cells in CA1, CA3 and the pre-subiculum).

A key question, therefore, is whether analysis approaches that are sensitive to submaximal patterns of activity, as is Multivariate Pattern Analysis (MVPA) which can identify distinct patterns in fine-grained activity responses to different stimulus categories, would be more sensitive to potential between-category differences outside of the subicular complex (for reviews of this method see Haxby, 2012; Haxby et al., 2014; Norman et al., 2006; Weaverdyck et al., 2020; Yang et al., 2012). However, this approach has not been applied in the context of hippocampal category selective effects in complex perception.

Additionally, although much recent work has placed scenes at the heart of the hippocampal contribution to cognition (Zeidman & Maguire, 2016; Murray, Wise and Graham, 2018), the spatially modulated hippocampal circuit involving anteromedial subicular complex may be completed by one including lateral hippocampus (corresponding with the location of the prosubiculum/CA1). Unlike the putative spatially modulated circuit, this is proposed to carry face/object information (Dalton & Maguire, 2017, Dalton et al., 2018) due to direct links with the perirhinal cortex (PRC; Insausti and Muñoz, 2001), which is critical to performance of face oddity judgement tasks (Lee et al., 2005; Berhmann et al., 2016). These parallel scene and face/object processing streams are also apparent in the entorhinal cortex; human functional connectivity work has shown PRC- and PHC-preferential functional connectivity with anterior-lateral and posterior-medial entorhinal cortex, respectively which, in turn, show different connectivities with lateral subicular complex (and CA1 border) and medial subicular complex (Grande et al., 2022; Maass et al., 2015). It may be possible, therefore, that more sensitive approaches (e.g., MVPA) can identify patterns of activity that are sensitive to face-related information during visual perception, particularly in more lateral parts of the hippocampal formation, thus revealing two parallel processing streams for different information categories, converging in different locations of the hippocampus.

Here, then, we addressed two specific questions. First, the extent to which, during perceptual discrimination, hippocampal activity patterns specific to scenes versus other visual categories are focused on a putative anteromedial subiculum ‘hub’ (potentially corresponding with the location of distal subiculum or pre/para-subiculum) or are distributed more widely throughout the hippocampus. Second, whether, as suggested by Dalton & Maguire (2017) patterns of activity within the lateral hippocampus (corresponding with the location of the pro-subiculum/CA1) will carry information relevant to the visual discrimination of faces/objects.

To address these questions, we applied MVPA to the high-resolution fMRI data of our original 7T oddity perceptual discrimination study (Hodgetts et al., 2017). Specifically, we used support vector machine (SVM) searchlights to examine whether (a) scene trials could be distinguished from face, object, and shape-size trials, and (b) face trials could be distinguished from scene, face, and shape-size trials, based on the activity patterns across both the whole hippocampus, as well as the extended fMRI field-of-view (FOV), that encompassed regions of retrosplenial cortex (RSC) and parahippocampal gyrus (PHC) considered to form a ‘core’ scene processing network (Baldassano et al., 2016; Hodgetts et al., 2016; Epstein and Baker, 2019), and regions of inferior- and superior-temporal regions considered to form a ‘core’ face network (Haxby et al., 2000; Bernstein & Yovel, 2015; Grill-Spector et al., 2017). Participants’ hippocampal subfields were also manually segmented on ultra-high-resolution images and co-registered to a standard template. This generated a novel probabilistic atlas of hippocampal subfields that allowed us to interrogate the location and extent of scene- and object/face-related category information based on classical definitions.

Initially, searchlight classification was carried out within the hippocampus only, to the unsmoothed fMRI data (1.2 mm isotropic voxels) at the individual-level. Uncorrected significance maps (accuracy significantly differing from zero) were then overlaid in MNI space to provide insight into both the spatial distribution and inter-individual variability of scene (and object/face) related information within the hippocampus (building on Hodgetts et al., 2016). Thus, as well as potentially increasing sensitivity to the information contained within hippocampal subfields (Weaverdyck et al., 2020), this more exploratory searchlight approach has the potential to identify the distribution of informational content both within and across classic subfield ROI boundaries. Accordingly, we predicted gradients of scene and face classification overlap would differ across the hippocampus and within the subicular complex and CA1, such that scene classification overlap would be greater more medially, while face classification overlap would be greater more laterally. Second, searchlight classification was applied across the entire FOV at the individual level, but accuracy maps were transformed to MNI space where conservative, multiple comparisons corrected statistical analyses were performed. We predicted that scene selective regions would overlap with the anteromedial subicular complex, CA1 and cortical scene sensitive regions, the RSC and parahippocampal gyrus. Conversely, we predicted that face sensitive regions would overlap with perirhinal cortex and cortical face sensitive regions such as the fusiform cortex and posterior superior temporal sulcus.

## 2. Methods

This work is a secondary analysis of previously published data (Hodgetts et al., 2017). Despite some aspects being the same, for completeness we describe the methods in full here.

### 2.1. Subjects

Twenty-five participants (healthy with no history of neurological or psychiatric illness; 16 females; 9 males; age range = 18-35 years; age mean = 25; age SD = 4) were recruited from the University of Oxford and Oxford Brookes University. They were fluent English speakers with normal/ corrected-to-normal vision. The research was approved by the University of Oxford Central University Research Ethics Committee and the Medical Sciences Interdisciplinary Research Ethics Committee, and each participant provided written informed consent before the experiment.

### 2.2 MRI data acquisition

A Siemens 7T Magnetom scanner, with a 32-channel head coil (Nova Medical, MA), was used to acquire MRI data. Whole-head T1-weighted data were produced with a MPRAGE sequence (1 x 1 x 1 mm; TE = 2.82 ms; TR = 2200 ms; flip angle = 7°). Blood-oxygen level-dependent (BOLD) data were acquired using a T2*-weighted echo planar imaging (EPI) sequence. The three oddity task fMRI runs consisted of 212 volumes and took approximately 7 minutes each (voxel size = 1.2 x 1.2 x 1.2 mm; slices = 30; TE = 25 ms, TR = 2000 ms; flip angle = 90°; partial FOV = 192 mm (Figure. 1C); partial Fourier = 6/8; parallel imaging with GRAPPA factor = 2; bandwidth = 1562 Hz/Px; echo spacing = 0.72 ms). Slices were oriented parallel to the hippocampal long axis. Slice acquisition occurred in a descending odd-even/interleaved order. To allow for magnetization equilibrium, three volumes were discarded at the beginning of each run. A whole brain T2*-weighted EPI volume was also acquired using identical image parameters, to facilitate co-registration of partial FOV images. To improve registration and reduce image distortion from magnetic-field inhomogeneity, a field map was acquired with the same slice orientation as the functional acquisition (TE 1 = 4.08 ms; TE 2 = 5.1 ms; TR = 620 ms; FOV = 192 mm; flip angle = 39°). Two T2*-weighted ultra-high-resolution structural images were acquired with opposite phase encoding directions (left-to-right; right-to-left; voxel size = 0.6 x 0.6 x 0.6 mm; slices = 44; TE = 25.7 ms; TR = 50 ms; partial Fourier = 6/8; FOV = 192 mm). The experimenters and radiographer aligned slices orthogonal to the hippocampal main axis, by visual inspection.

### 2.3. MRI pre-processing

FMRIB Software Library (FSL) (Jenkinson et al., 2012) was used to process the fMRI data. The raw data were first converted to NIfTI format. The T1-weighted images were then stripped of non-brain tissue using BET (Smith, 2002). Bias field correction was carried out using the Enhancing Neuro Imaging Genetics through Meta Analysis (ENIGMA) protocol pipeline (Thompson et al., 2020). Prior to analysis, each functional run was motion corrected and co-aligned (registered to the middle volume of the second run) using MCFLIRT (Jenkinson et al., 2002). Additional EPI pre-processing was carried out using the FMRI Expert Analysis Tool (FEAT) Version 6, including high-pass temporal filtering (Gaussian-weighted least-squares straight line fitting, with σ = 50 s) and field map unwarping using FUGUE (Jenkinson et al., 2002). No spatial smoothing was applied. Time-series statistical analysis was carried out using FMRIB’s Improved Linear Model (FILM) with local autocorrelation correction (Woolrich et al., 2001).

#### 2.3.1. MRI data exclusion

Our *a priori* threshold for participant exclusion based on motion was one EPI voxel (1.2 mm): no participant exceeded this threshold. Two participants were removed as one had an incidental finding on their MRI, and another had excessive susceptibility artefacts in the temporal lobe which impeded hippocampal segmentation. Therefore, data from 23 subjects were included.

### 2.4. Hippocampal subfield segmentation and co-registration

To contextualise the MVPA searchlight results, we created probabilistic hippocampal subfield ROIs (CA1, CA2/3, dentate gyrus, and subiculum) based on manual segmentations within our participants. Hippocampal subfields were initially manually segmented (by C.J.H.) on participants’ ultra-high-resolution T2*-weighted images using a 7T-specific protocol based on Wisse et al. (2012), and then co-registered to each individual’s fMRI space (see Hodgetts et al., 2017). These individual-subject ROIs were then co-registered to the 1mm MNI template, binarized, summed, and (to aid interpretation and visualization) thresholded so that each ROI covered voxels included in at least 25% of participants (Figure 1D; see Syversen et al., 2021 for an example of use of this threshold). The full (unthresholded) probabilistic ROIs are freely available at: https://osf.io/xc4wa/.

### 2.5 The oddity judgement task

Participants completed a simultaneous visual discrimination ‘oddity’ judgement task (Buckley et al., 2001; Lee et al., 2005). On a given trial, subjects were presented with three stimuli (top centre; bottom left; bottom right) and asked to choose the odd-one-out as quickly and as accurately as possible. In each trial, the triplets of images were presented on a white background (Figure 2A). The scene stimuli were greyscale photographs of real outdoor locations (in Cardiff City Centre) and were unfamiliar to participants (see Shine et al., 2015). Two scenes depicted a single location from different viewpoints and one scene depicted a different, but highly similar, location. Face stimuli were greyscale photographs of human faces (half male and half female) and were obtained from the Psychological Image Collection at Stirling (PICS, http://pics.stir.ac.uk/). Individual faces were overlaid on a black frame of 170 x 216 pixels. Two faces were the same individual at different viewpoints, or with a different facial expression, and the third (odd) image was a different face, of the same sex, presented from a different viewpoint. Objects were obtained from the Hemera Photo-Objects 50,000, Volumes 1-3 (Hemera Technologies, Quebec). The trials included two identical objects presented from different viewpoints, and the third object was different but from the same subordinate-level object category. For the ‘size’ trials, three black squares were shown. Two of which were identical in size and a third square was either slightly larger or smaller; the difference in length between target and non-targets could vary between 9 and 15 pixels. The position of the squares was jittered so that none of the edges lined up along vertical or horizontal axes. All stimuli were trial-unique (i.e., each stimulus was shown only once). Immediately prior to scanning, participants were shown a practice trial for each category (not repeated during the experiment) and indicated to the experimenter their correct response.

**Figure 2.**
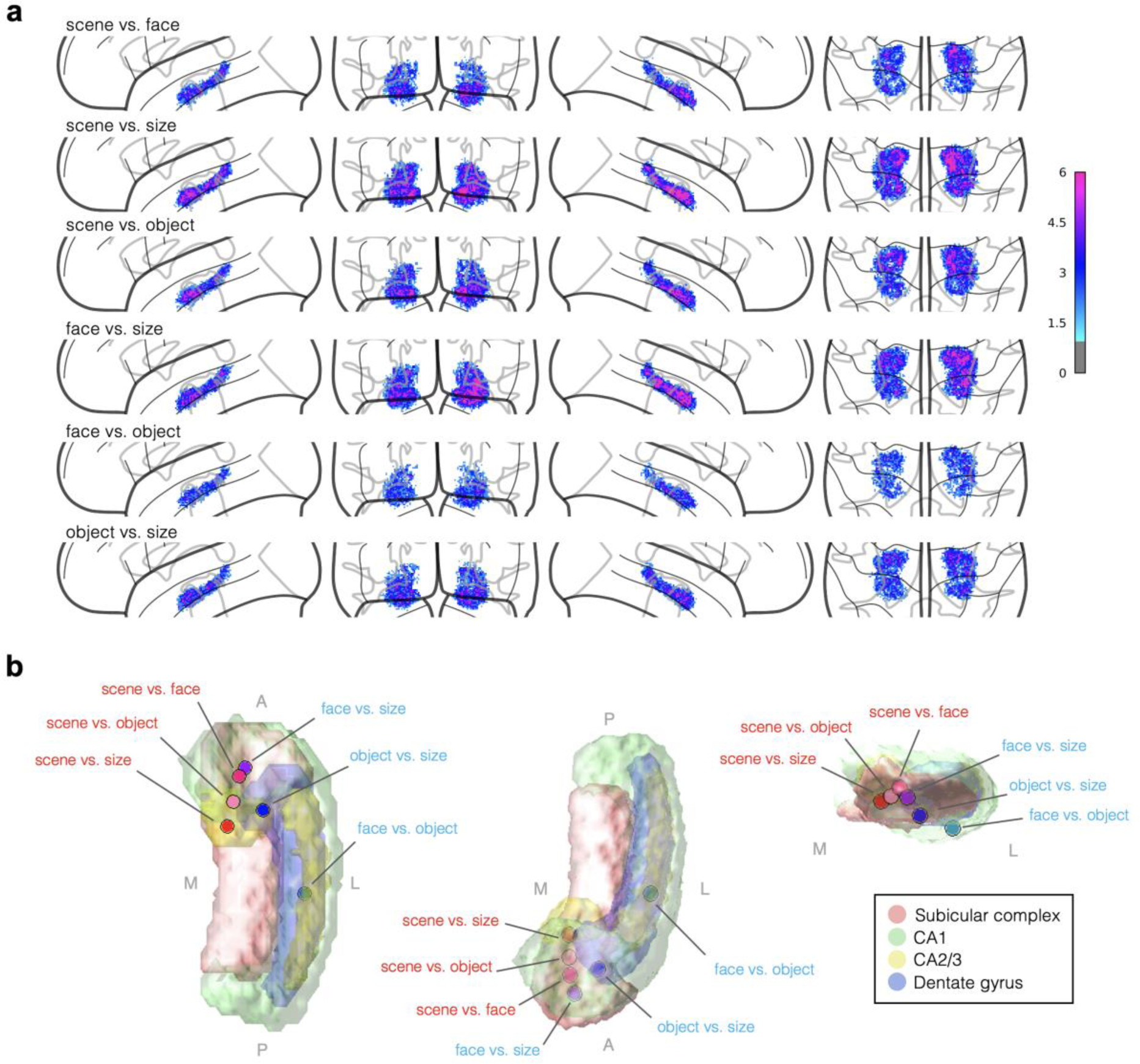
Overlap statistical hippocampal searchlight maps. A) Overlap maps constructed by summing participant significance maps from individual level Binomial statistical tests against chance level. Maps have a lower threshold of 1, pink colours indicate higher overlap than blue colours. B) Peaks of the overlap maps plotted on three view angles of the right hippocampal subfields (the peaks and ROIs are coloured according to the key to the right).

The MRI task was projected onto the screen behind the subject using an Eiki LC-XL100 projector system (Resolution: 1024 x 768; Refresh Rate: 60Hz). The paradigm was coded using Presentation (Neurobehavioural Systems, CA), and button presses were recorded with a right-hand MR compatible button box. Trials were presented for 5500 ms with a jittered inter-trial interval (blank screen) of 500-2500 ms (Figure 2B). The task was carried out over three fMRI runs (each run was ∼7 minutes in duration). Trials were presented in mini-blocks of three consecutive trials of the same condition (either three scenes, faces, objects, or sizes). The order of which was counterbalanced across participants. 15 trials of each stimulus category were presented, per run, totalling to 45 trials per condition. Each condition had the same number of targets (odd-one-out) appearing in each triplet position (i.e., top centre; bottom left; bottom right).

### 2.6. General linear model

Double-gamma hemodynamic response functions were used to model BOLD signals. Each trial, per run, was entered as an explanatory variable (EV) within a General Linear Model (GLM), with event durations of 5.5 s, using FEAT (Woolrich et al., 2001). This resulted in three first-level models (for each of the runs), each containing 60 main EVs. (15 per condition: scene, object, face, size) and a motor regressor corresponding to button press onset on each oddity task (duration = 0 s). An additional confound matrix was added to the GLMs to account for volume-wise non-linear motion effects using FSL Motion Outliers. This resulted in 180 t-statistic images for each participant, derived from the parameter estimates. The t-statistic images were then used in subsequent MVPA searchlight analyses.

### 2.7. Searchlight analysis

Linear support vector machine (SVM) classifier-based searchlight analyses (chosen because it’s resilience to overfitting) were conducted using the MVPA-light toolbox (Treder, 2020) in MATLAB R2015A. Neighbours were defined as 3×3×3 cubes of voxels. Considering the high computational load when searching high resolution data, we used five k-folds (45-9 trials used as the test set), with two repeats (a different test set randomly assigned in each repeat, and classifier accuracy values averaged across repeats). Searchlight analyses were carried out in native space and resulting accuracy maps were then warped to the T1-weighted 1mm MNI standard template (skull stripped) using FNIRT (Jenkinson et al., 2012). We performed two types of searchlight analyses, the first of which was restricted to the hippocampus, and designed to be sensitive to category information, and to allow accommodation of individual differences in intra-hippocampal locations of category selectivity. The second included data from the entire FOV and incorporated traditional conservative group-level statistics, designed to combat the potential bias (predisposition to produce false positives) of the first analysis.

#### 2.7.1. Hippocampal searchlight

First, 2-way classification searchlights of all stimulus category pairs were performed within the hippocampus of each individual subject. As individual variability in the location of category-selective clusters may cause informative clusters to be missed at the group-level (Etzel et al., 2013), we sought to conduct statistical inference at the individual level, prior to creating probabilistic overlap maps on our novel hippocampus MNI template (see Hodgetts et al., 2016; Kaplan and Meyer, 2012; for similar approaches). Statistics were performed at the individual level by carrying out binomial tests on the individual searchlight accuracy result maps (for each 2-way classification) assuming a chance level of 50% (Treder, 2020). Masks for each participant were constructed by selecting voxels where p-values < 0.05. These individual-level masks were then warped to template space using FNIRT, binarised, and summed across the 23 individuals. Therefore, this resulted in 6 searchlight contrast overlap maps (scene vs. face, scene vs. size, scene vs. object, face vs. size, face v object, object vs. size). We also attempted to contrast locations of scene and face information specifically by further combining these overlap maps into scene and face ‘hotspot’ maps. The scene hotspot map was constructed by summing scene vs. face, scene vs. size, scene vs. object overlap maps, and subtracting face vs. size, face vs. object overlap maps. The face hotspot map was constructed by summing scene vs. face, face vs. size, face vs. object overlap maps and subtracting scene vs. size, scene vs. object overlap maps. Note, the focus on faces in the current investigation (as opposed to faces *and* objects) is based on previous evidence showing that hippocampal lesions have stronger influence on face discrimination performance using similar oddity tasks (Behrmann et al., 2016; Chang et al., 2023).

#### 2.7.2. Whole FOV searchlight

We also sought to examine scene and face selectivity within our extended FOV (e.g., in posteromedial and parahippocampal cortex) using traditional conservative group-level statistics. To do this we applied permutation tests to group-level searchlight data. First, native space accuracy maps of each category pair (2-way classification) were warped to standard MNI space and combined into a 4D file. Second, 0.5 was subtracted from each map so that chance level was zero. A threshold of −0.2 was applied to that regions that were zero remained zero, rather than becoming −0.5 (−0.2 was arbitrarily chosen after inspections of image histograms showed a curtailing of voxels with accuracy levels below this value. One-sample permutation tests were carried out using FSL’s ‘Randomise’ (Winkler et al., 2014). We used threshold-free cluster enhancement (TFCE) (Smith & Nichols, 2009) and 5000 permutations, and, as the FOV differed slightly between individuals, a mask that included voxels where all participants had values (Figure 1C). The resulting t-maps were masked using the corrected p-values maps and thresholded at p = 0.0083 (0.05 / 6 tests). To isolate scene and face information specifically, conjunction analyses were then performed, by combining significance masks from the searchlight contrasts (binary masks multiplied together). Specifically, a scene-selective map was the product of all the significance masks from searchlight contrasts that included scene trials (scene vs. face × scene vs. size × scene vs. object) with the product of all the significance masks from the non-scene searchlights (face vs. size × face vs. object × object vs. size) removed (and *vice versa* for face selective regions).

#### 2.7.3. Data sharing and open practices

Anonymised output data and the code used within this project are freely available at https://osf.io/4vgk9/, so that the figures and values reported in this manuscript are reproducible. However, ethical restrictions, relating to General Data Protection Regulation, do not allow for the public archiving of the raw study data. Access to pseudo-anonymized data could be granted after signing and approval of data-transfer agreements. For this, readers should contact Dr Carl Hodgetts.

## 3. Results

### 3.1. Oddity judgement task behavioural results

Participants achieved high and matched decision accuracies across image categories. The mean proportion correct scores (and SDs) of the scene, face, object, and size conditions were: 0.76 (14), 0.76 (14), 0.77 (16), 0.76 (15), respectively. Detailed analyses on behavioural performance are available in our previous publication (Hodgetts et al., 2017).

### 3.2. Searchlight MVPA results

#### 3.2.1 Hippocampal searchlight classification

First, we examined the spatial distribution and inter-individual variability of scene and non-scene-selective information within our hippocampal ROI. As noted in the Methods, we conducted a two-category SVM searchlight within each individual’s hippocampal ROI for each of the discrimination pairs (scene vs. face, etc), then co-registered the resulting (binarized) maps of significant classification voxels (> 50%) to the standard template before summing the maps across subjects to create overlap maps. The mean number of voxels (and SDs) reaching significance in the binomial tests for each hippocampal searchlight contrast were: scene vs. face: 870 (511); scene vs. size: 1288 (889); scene vs. object: 1047 (402); face vs. size: 1236 (670); face vs. object: 642 (267); object vs. size: 891 (494).

Figure 2A shows the overlap for each of the SVM hippocampal searchlights (greater values equals larger overlap between subjects). As can be seen, category selectivity appeared to cover most of the hippocampi ROIs, but the highest overlap between participants was generally seen anteriorly (except for the face vs. object overlap map), and in the right hippocampus. When examining the location and distribution of these overlap peaks (voxel locations where most participants had significant voxels) with respect to our probabilistic subfield ROIs (Figure 2B), we found that the scene-including classification overlap maps had the highest overlaps medially, while the highest overlaps of the face/object-including classification overlap maps were more lateral. The scene vs. size overlap peak was the most medial, located in the anteromedial subicular complex, close to the superior subicular complex/CA1/CA23 ROI borders (maximum overlap between individuals = 10; x = 18, y = −17, z = −18, 43% subicular complex, 20% CA1, 20% CA23, 3% DG). The scene vs. object overlap peak was located at the superior subicular complex/CA1 border (7; x = 19, y = −14, z = −20; 52% subicular complex, 36% CA1, 4% CA23, 2% DG). The scene vs. face classification overlap map had the highest overlap in anterior CA1 (8; x = 20, y = −11, z = −21; 67% CA1, 34% subicular complex, <1% DG).

With regards to our face/object classification overlap maps, we found that the face vs. object classification overlap peak was the most lateral and posterior, located in CA1 (6; x = 31, y = −25, z = −15; 86% CA1, 4% DG). The object vs. size classification overlap map had the highest overlap in the subiculum, but this was more lateral and inferior than the scene classification overlap map peaks, and situated close to the subiculum’s lateral border with CA1 (7; x = 24, y = −15, z = −23; 66% subicular complex, 9% CA1, 3% DG). The face vs. size classification overlap map had the highest overlap in the anterior subiculum, close to its superior border with CA1 (8; x = 21, y = −10, z = −24; 63% subicular complex, 28% CA1).

With the aim of comparing where selective scene and face information is most likely to be located, further ‘hotspot’ overlap maps (Figure 3C) were constructed by 1) summing together hippocampal searchlight overlap maps from comparisons including scenes, and subtracting the sum of overlap maps from comparisons including faces (‘scene hotspot’), and 2) summing together hippocampal searchlight results from comparisons including faces, subtracting the sum of overlap maps from comparisons including scenes (‘face hotspot’; see Methods). Again, a medial-lateral gradient was apparent, with the scene hotspot map showing higher values medially, and the face hotspot map showing higher values laterally. The maximum value for the scene hotspot maps was in left medial inferior CA1 and superior subiculum boarder (x = −17, y = −15, z = −19; 45% CA1, 39% subicular complex). The maximum value for the face hotspot maps was located in right lateral hippocampus, in the inferior border of the DG with CA1 (x = 32, y = −23, z = −14; 70% DG, 29% CA1; Figure 3). These peak locations remained when the hotspot maps were restricted to the CA1 ROI (Euclidean distance between scene and face hotspot maps maximum values = 50mm).

**Figure 3.**
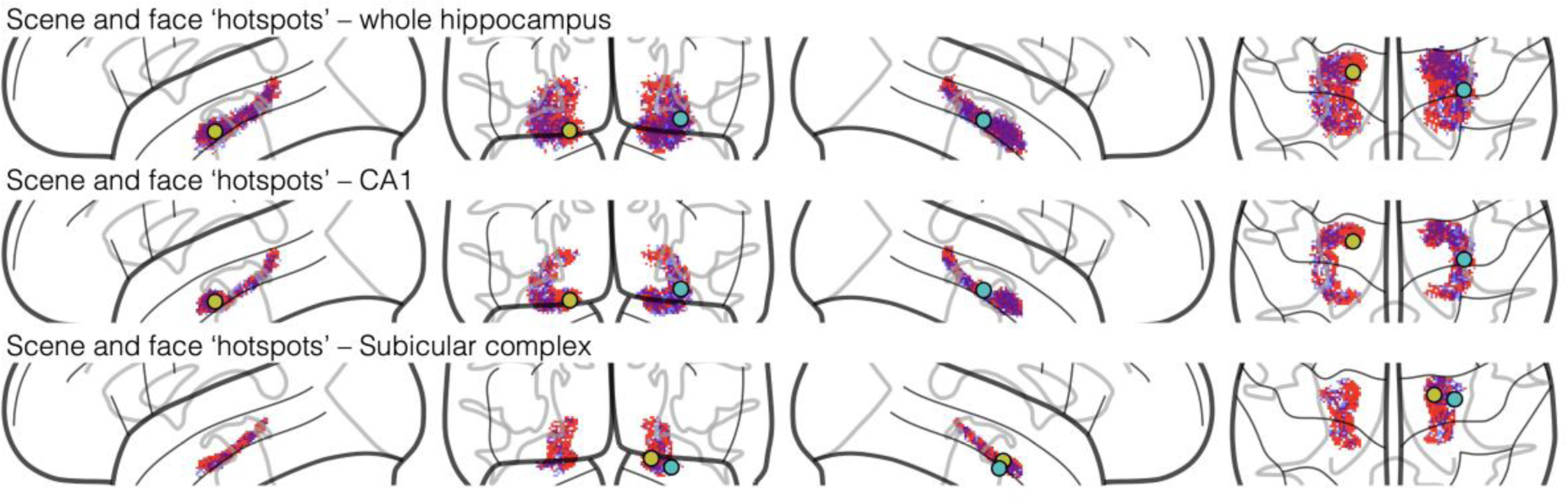
Overlap statistical hippocampal scene and face ‘hotspot’ searchlight maps. ‘Hotspot’ maps constructed by summing together searchlight results from comparisons including scenes, with those results from comparisons including faces subtracted (reds) and constructed by summing together searchlight results from comparisons including faces, with those results from comparisons including scenes subtracted (blues). Overlaid dots represent the locations of maximum overlap for the scene ‘hotspot’ map (yellow dot) and the face ‘hotspot’ map (light blue dot). Note that an arbitrary minimum threshold of 3 was applied to increase clarity in the image.

This medial-lateral pattern was still apparent when the hotspot maps were restricted to the subicular complex ROI. Maximum values for the scene and face hotspot maps were located in right anterior medial (x = 19.0, y = −14.0, z = −20.0; 52% subiculum, 36% CA1, 4% CA2/3, 2% DG) and lateral areas, respectively (x = 28.0, y = −16.0, z = −24.0; 22% subiculum, 21% CA1; Euclidean distance between subiculum scene and face hotspot maps maximum values = 10mm).

#### 3.2.2 Whole FOV searchlight classification

To examine the whole fMRI FOV, we adopted a more conservative, group statistical approach (group-level nonparametric permutation inference). Specifically, we initially conducted SVM searchlights across each individual’s FOV for each discrimination of interest. Accuracy maps were then co-registered to the standard MNI template before Randomise tests (cluster permutation tests against chance level) were run. Each 1 vs. 1 searchlight classification (scene vs. face, scene vs. object, scene vs. size, face vs. size, face vs. object, object vs. size) resulted in significant clusters.

To isolate regions selective to scenes and not faces within the ‘scene vs. face’ significance mask, significant voxels from ‘face vs. size’ were removed (Figure 4A). Then, to isolate regions selective to faces and not scenes within the ‘scene vs. face’ significance mask, significant voxels from scene vs. size’ were removed (Figure 4B). Scene regions were found in right anteromedial subicular complex (maximum ROI probability: 64%, x = 19 y = −18 z = −20; 25 voxels above 25%) and bilateral lateral CA1 (maximum ROI probability: 67%, x = 31 y = −31 z = −12; 229 voxels above 25%). Face regions were also apparent in left anteromedial subicular complex (maximum ROI probability: 29%, x = −17 y = −30 z = −13; 2 voxels above 25%) and lateral CA1 (maximum ROI probability: 68%, x = 30 y = −32 z = −11; 78 voxels above 25%), albeit to a lesser extent.

**Figure 4.**
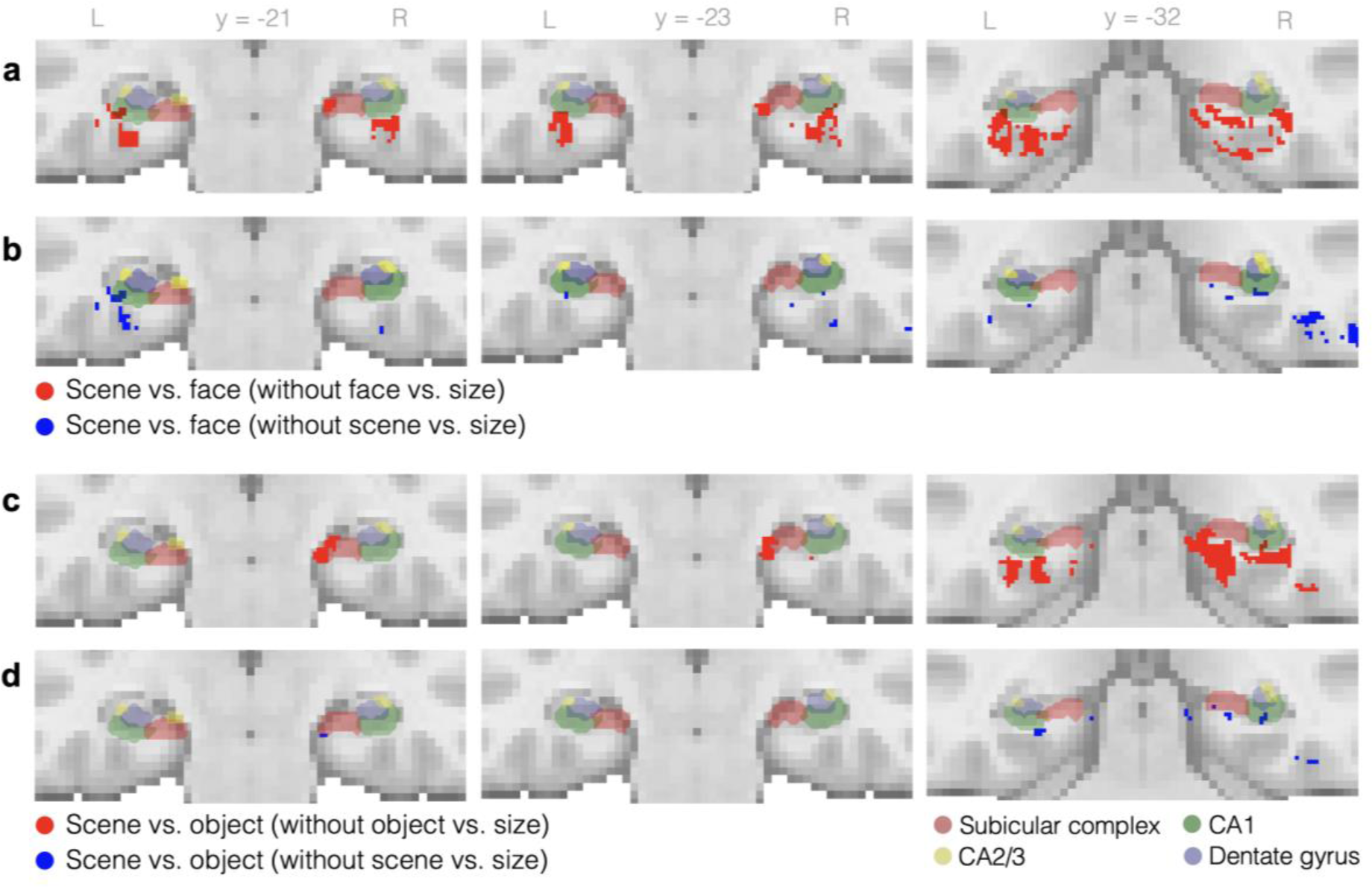
Scene vs. Face and Scene vs. Object Conjunction maps. A) Scene regions from the scene vs. face searchlight map (’face vs. size’ removed). B) Face regions from the scene vs. face searchlight map (‘scene vs. size’ removed). C) Scene regions from the scene vs. object searchlight map (‘object vs. size’ removed). D) Object regions from the scene vs. object searchlight map (‘object vs. size’ removed). The resulting maps are overlaid onto our group subfield ROIs. DG: Dentate Gyrus, L: Left, R: Right. Image created using Nilearn (Abraham et al., 2014) for Python.

The same was carried out for ‘scene vs. object’ (Figure 4C-D). Again, a scene region was found in right anteromedial subiculum (maximum ROI probability: 75%, x = 20 y = −18 z = −19; 156 voxels above 25%), but CA1 regions were more medial than those in the scene vs. face mask (maximum ROI probability: 78%, x = 30 y = −31 z = −11; 117 voxels above 25%). In addition, the scene region overlapped with medial CA23 (maximum ROI probability: 59%, x = 20 y = −18 z = −16; 70 voxels above 25%). Object regions were also apparent in anteromedial subicular complex (maximum ROI probability: 75%, x = 20 y = −18 z = −16; 33 voxels above 25%), CA1 (maximum ROI probability: 78%, x = 30 y = −31 z = −11; 29 voxels above 25%) and CA23 (maximum ROI probability: 52%, x = 19 y = −17 z = −13; 14 voxels above 25%), to lesser extents.

Lastly a purely **scene-selective map** (Figure 5-6) was defined as the conjunction of significant voxels from all scene-discriminating searchlights (‘scene vs. face’, ‘scene vs. size’ and ‘scene vs. object’), with the conjunction of significant voxels from all non-scene-discriminating searchlights (‘face vs. size’, ‘face vs. object’ and ‘object vs. size’) removed. Scene-selective regions were found to overlap with the anteromedial portion of the right subicular complex ROI (maximum ROI probability: 55%, x = 19 y = −21 z = −21; 29 voxels above 25%), a region we interpret as corresponding to distal subiculum/presubiculum/parasubiculum (Figure 5A), and left posterior lateral CA1 (maximum ROI probability: 50%, x = −33 y = −34 z = −10; 31 voxels above 25%), a region we interpret to correspond to posterior proximal CA1 (Figure 5A). Supporting our interpretations, Figure 5B shows the locations of these scene-selective map ROI maximum probabilities in relation 1) to the anterior-posterior hippocampus border (defined as the uncal apex at y = −22; as defined by Poppenk et al., 2013; Zeidman and Maguire, 2016), 2) a previously defined anteromedial hippocampus location (used by Zeidman, Mullally, et al., 2015; Zeidman and Maguire 2016 as a seed region to explore anteromedial hippocampus connectivity, defined through the conjunction of activation during recalling and imagining scenes), and 3) a previously observed peak BOLD location within the anteromedial hippocampus during a scene construction task, interpreted as residing in pre/parasubiculum (Dalton et al., 2018). In addition, the scene-selective map overlapped with known scene selective areas, the posterior PHC and RSC (Figure 6A). The three most overlapping regions within FSL’s Harvard-Oxford cortical structural atlas were: precuneus cortex (16% overlap), superior lateral occipital cortex (11% overlap); and lingual gyrus (10%).

**Figure 5.**
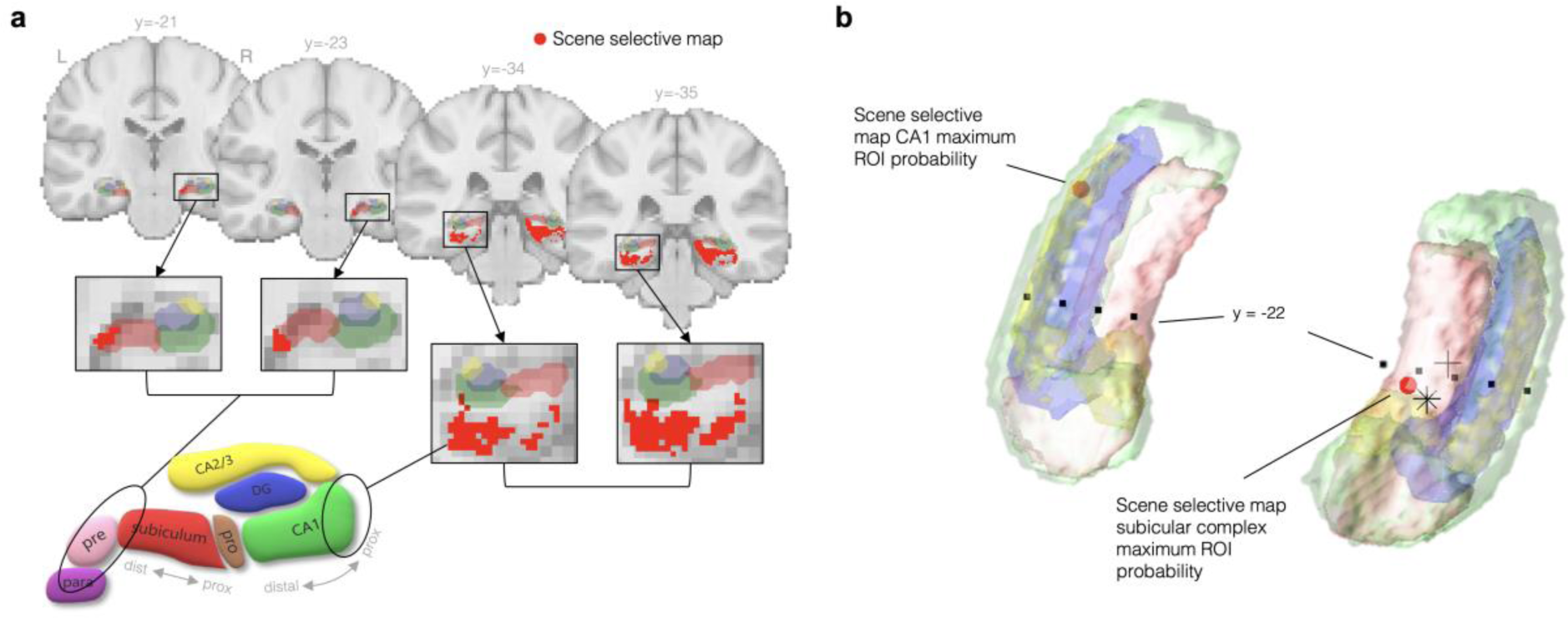
The scene-selective and face selective conjunctive maps within the hippocampus. A) The ‘scene-selective’ map plotted onto multiple coronal template slices. The four coronal ‘zoomed-in’ images, show inclusion of the right anteromedial subicular complex and inclusion of lateral CA1 regions. MNI slice coordinates are stated by each image. Our subfield ROIs were overlaid onto this map, and their colours correspond to the drawn subfield diagram. The subfield diagram illustrates our interpretation of the results and includes subfield delineations not possible with our current methods. The circles indicate that we consider the portion of the scene-selective map that overlaps with the medial subicular complex to incorporate distal subiculum/presubiculum/parasubiculum areas. We also consider the portion of the scene-selective map that overlaps with lateral CA1 to incorporate part of the proximal aspect of CA1. B) The locations of these scene-selective map ROI maximum probabilities in relation to the anterior-posterior hippocampus border (y = −22; Zeidman and Maguire (2016). To contextualise these findings, a previously defined anteromedial hippocampus location (Zeidman, Mullally, et al., 2015; Zeidman and Maguire 2016), and a previously found anteromedial hippocampus scene construction peak within the pre/parasubiculum (Dalton et al., 2018) is also shown. DG: Dentate Gyrus, L: Left, para: parasubiculum, pre: presubiculum, pro: prosubiculum, R: Right. Images created using Nilearn (Abraham et al., 2014) for Python, MATLAB, and Microsoft products.

**Figure 6.**
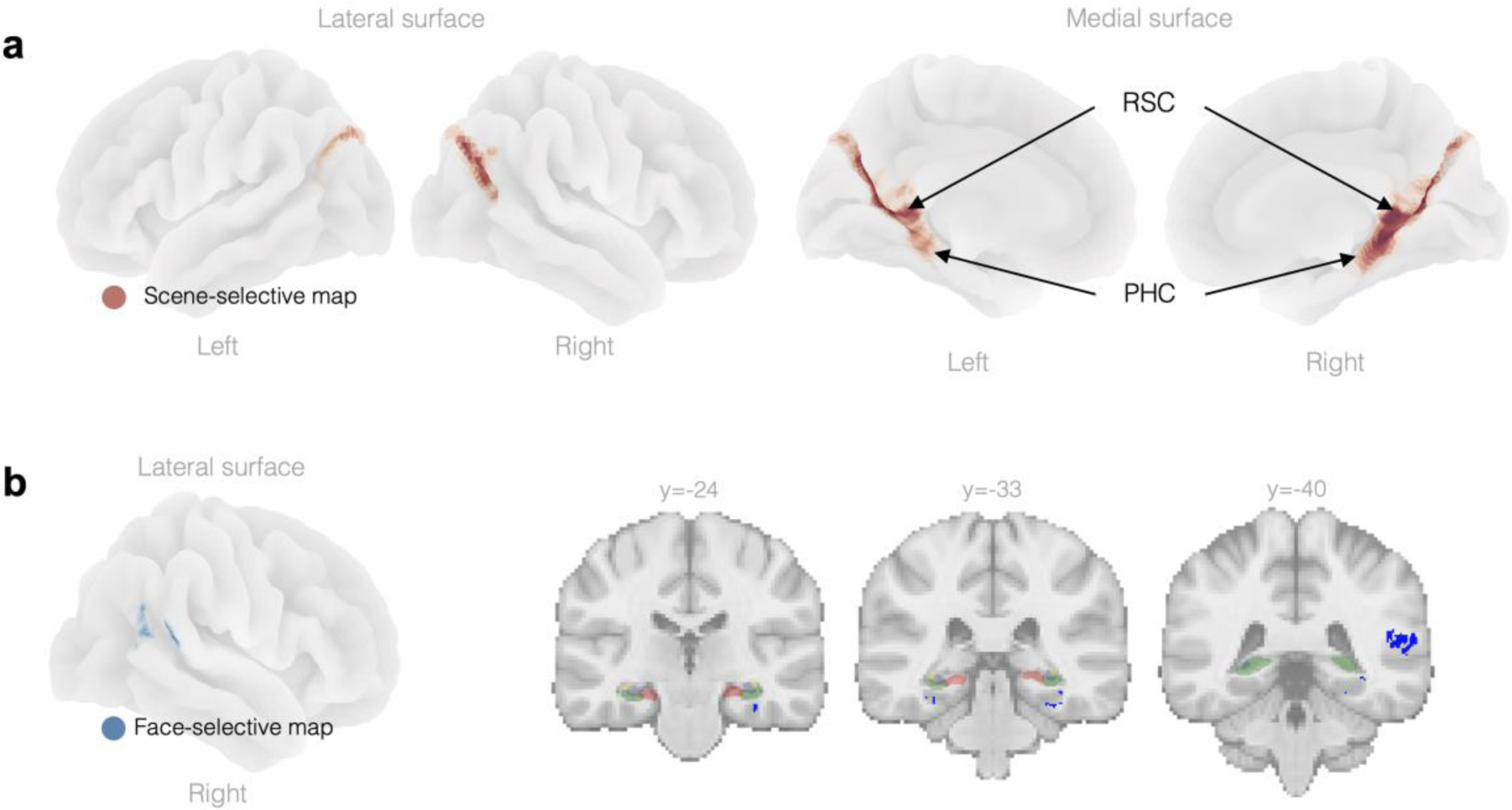
The scene-selective and face selective conjunctive maps within the whole fMRI FOV. A) The ‘scene-selective’ map plotted onto template brain surfaces, which shows the scene-selective map to overlap with areas of the superior occipital, PHC and RSC regions. D) The ‘face-selective’ map plotted onto a template brain surface (left) and onto multiple coronal slices (right). PHC: Parahippocampal cortex, RSC: Retrosplenial Cortex. Images created using Nilearn (Abraham et al., 2014) for Python, MATLAB, and Microsoft products.

In contrast, a **face-selective map** (Figure 6B) was defined as the conjunction of significant voxels from all face-discriminating searchlights (‘scene vs. face’, ‘face vs. size’ and ‘face vs. object’), with the conjunction of significant voxels from all non-face-discriminating searchlights (‘scene vs. size’, ‘scene vs. object’ and ‘object vs. size’) removed. This face-selective map was right lateralised and did not overlap with our hippocampal ROIs. The three most overlapping regions within FSL’s Harvard-Oxford cortical structural atlas were: angular gyrus (18% overlap); posterior supramarginal gyrus (12% overlap); and temporooccipital gyrus (13% overlap), and there was also overlap with the temporal fusiform cortex (4% overlap).

## 4. Discussion

In the current study, we advanced our knowledge of hippocampal contributions to high-level scene perceptual discrimination by: 1) Using MVPA to improve sensitivity to fine-grained category-selectivity patterns of hippocampal BOLD activity, thereby revealing decodable scene information outside of the subiculum (previously identified as a hippocampal ‘hub’ of scene information based on magnitude of univariate BOLD activity); 2) using a subfield agnostic searchlight approach, thereby revealing precise locations of category selectivity that do not necessarily conform to traditional anatomical subfield boundaries and; 3) using both subject-level and group-based analysis in order to account for the different biases present in each (see Etzel et al., 2013), which helped to reveal a transverse and longitudinal gradient of scene category selectivity in the subiculum and CA1. In sum, this approach allow us to map a scene representation pathway in humans, which included regions within the anteromedial (distal) subicular complex and lateral (proximal) CA1, alongside cortical regions that form part of a parieto-medial temporal pathway (Kravitz et al., 2011) implicated elsewhere in spatial navigation and scene-based cognition, namely the posterior PHC and RSC (Boccia et al., 2016, Soto et al., 2012, Margulies et al., 2009, Goodale & Milner, 1992; Epstein & Baker, 2019). In addition, we provide preliminary evidence suggesting that areas within the lateral hippocampus carry information relevant to discriminating faces. We discuss the implications of these findings for accounts of hippocampal contributions to higher-level perception in the following sections.

### 4.1. Subject-level analysis indicates gradients of hippocampal category sensitivity

Both the ‘overlap’ and ‘hotspot’ maps (see Figures 2 & 3) demonstrated a gradient-like pattern of scene selectivity, with scene information being preferentially processed anteriorly on the longitudinal (i.e., anterior - posterior) axis and medially along the transverse (i.e., distal – proximal/ medial – lateral^1^) axis. In contrast, face and object information showed a somewhat more diffuse pattern, although there was evidence for increasing face selectivity laterally and anteriorly (see further discussion below). Mapping the data back to individual subfields, we additionally show that the same longitudinal and transverse gradients were present in both the subiculum complex and, notably, CA1. In addition, we report that the subiculum complex carried less face-sensitive information, with most of the face-selective voxels being present along the lateral subiculum border, whereas CA1 contained voxels relevant to the processing of both scenes and faces, with the aforementioned transverse gradient and a less pronounced longitudinal one.

Previous research has demonstrated that both functional and structural connectivity gradients in the hippocampus do not respect classical subfield boundary definitions (Aggleton & Christiansen, 2015; Dalton et al., 2022; Dalton et al., 2019). For scene selectivity, this aligns with current understanding of anterior-posterior hippocampal roles. The scene oddity task required participants to process whole scenes from multiple view-points, and the anterior hippocampus, in general, is proposed to bring together distributed visuospatial, object, and semantic information to construct spatially coherent internal representations of scenes (Sekeres et al., 2018; Zeidman & Maguire, 2016), and to process more global representations of space, relative to the posterior hippocampus (Evensmoen et al., 2013; McCormick et al., 2021: Nadel et al., 2013; Zeidman, Mullally, et al., 2015) (but see Ziegler et al., 2013 for a role for anterior hippocampus in fine-grained spatial discrimination), potentially as part of a wider role in ‘relational’ (Mayes et al. 2007; Olsen et al., 2012; Aly et al., 2013; Turk-Browne, 2019) or ‘compositional’ (Bakermans et al., 2023) processing. For face selectivity, our results demonstrating more lateral processing are in line with recent hippocampal structural connectivity research, which showed that cortical areas important for face processing (such as the temporal pole) preferentially terminate along the lateral borders (Dalton, 2022, Barton, 2022).

These results show the importance of studying longitudinal and transverse gradients across the hippocampus and within individual subfields. Further analysis of individual differences in these gradients may be useful in uncovering links to behaviours and pathology. Research on hippocampal connectivity has shown that individual differences in connectivity gradients are associated with phenotypes including mild cognitive impairment, Alzheimer’s disease, and more generalised differences in recollection memory performance (Borne et al., 2023, Przeździk et al., 2019). Larger scale investigations crossed with confirmatory gradient statistics (e.g., Przeździk et al., 2019) will be necessary to confirm the current findings in relation to scene and face processing.

### 4.2 Group-level analysis demonstrates a scene-processing pathway involving the distal subiculum and proximal CA1

At the group level, we combined significance masks across 2-category classification searchlights to identify scene, face, and object selective regions within ‘scene vs. face’ and ‘scene vs. object’ searchlight result maps and to create ‘scene-selective’ and ‘face-selective’ maps. Across all scene-discriminating searchlight combinations, regions within the anteromedial subicular complex and some inferior-distal voxels within CA1 were identified as scene-selective. The scene-selective map included the caudal inferior parietal cortex, bilateral anterior PHC, bilateral posterior RSC and extended into the medial temporal lobes, aligning with well-established scene networks (Baldassano et al., 2016; Epstein, 2008; Epstein & Baker, 2019; Hodgetts et al., 2016; Zeidman, Mullally, et al., 2015). In the hippocampus, this specifically included inferolateral edges of bilateral CA1, and an anteromedial portion of the right subicular complex (distal subicular complex). The part of the scene-selective map that invaded the subicular complex ROI also extended more medially, providing support for the suggestion, based on univariate fMRI analysis, that the pre/para-subiculum carries information important for scene processing including perceptual discrimination (Dalton & Maguire, 2017). Again, these results mirror recent evidence from structural connectivity analysis, which identified a similar anteromedial hub of hippocampal connectivity to cortical regions associated with scene processing (e.g., RSC) (Dalton et al., 2022).

Even with the use of high field (7T) fMRI, it is unclear whether our results show scene selectivity in the distal subiculum proper or pre/para-subiculum, or both (Dalton & Maguire, 2017; Ding, 2013). In terms of both its anatomical connectivity and cellular properties, the subicular complex appears perfectly situated to support scene processing as it is the primary source of hippocampal efferents (Aggleton, 2012; Bubb et al., 2017; Insausti & Munoz, 2001; Gigg, 2006) and, while many subiculum proper targets overlap with those of neighbouring CA1 (Matsumoto et al., 2019), almost all hippocampal projections to the anterior thalamic nuclei, mamillary bodies, and RSC originate in the subiculum proper (Aggleton & Christiansen, 2015; Frost et al., 2021; Gaffan, 1992; O’Mara & Aggleton, 2019). Structural connectivity between the subiculum and CA1 with the entorhinal cortex, along with converging connections to the RSC and PHC, have also been demonstrated (Kloosterman et al., 2003, Witter, 2006, Simonsen et al., 2022). Notably, recent functional connectivity analysis found that medial subiculum - entorhinal connectivity overlapped with entorhinal - PHC connectivity hotspots when participants were engaged in scene processing (Grande et al., 2022). These interconnected structures are key components of an extended hippocampal system thought to support spatial attributes of memory (Aggleton, 2012; see also Murray et al., 2017) as well as contributing to complex scene discrimination (Postans et al., 2014; Hodgetts et al., 2015). The subiculum proper, at least in rodents, also appears to contain several types of spatially-modulated neurons, including cells attuned to spatial position (Sharp, 2006), boundaries (Lever et al., 2009), corners (Sun et al., 2023) and direction of travel (Olson et al., 2017), as well as cells that seemingly fire conjunctively for several such properties needed to re-construct representations of spatial environments (Ledergerber et al., 2021; Sharma et al., 2022)

Conversely, while the subiculum proper is the main output region of the hippocampus, the pre/para-subiculum is primarily an input region, receiving input from areas including the subiculum proper, CA1, RSC and inferior parietal cortex (Honda & Shibata, 2017; Honda et al., 2022; Huang et al., 2021; Insausti & Munoz, 2001; Kravitz et al., 2011). A parieto-medial pathway originates in visuospatial regions of the parietal cortex, and connects to the parahippocampal cortex and RSC (Dalton & Maguire, 2017; Kravitz et al., 2011), two areas known to contribute to spatial processing (Cukur et al., 2016; Epstein et al., 2017; Hodgetts et al., 2016; Nasr et al., 2013), which in turn connect directly with the pre/para-subiculum (Dalton & Maguire, 2017). Likely related to these inputs, the pre/para-subiculum is known to contain grid, border and head-direction cells (Boccara et al., 2010; Robertson et al., 1999; Rolls, 2023), and indeed is considered a major componant of, and perhaps a site of visual information integration into, the head-direction system (Preston-Ferrer et al., 2016). Future fMRI studies at even higher spatial resolution, combined with directed functional connectivity measures (Reid et al., 2019), will be important in addressing how this network interacts during the processing of scenes.

The CA1 results are less easy to interpret, only existing at the edges of the ventral distal boundaries. This may simply reflect signal bleed from the PPA, which borders CA1, potentially caused by the spatial variability introduced either by the searchlights or from the registrations to standard space (Etzel, Zacks, & Braver, 2013). Alternatively, given previous literature indicating a role for CA1 in spatial processing, including place cells in rodents (e.g., see Brun et al., 2002; O’Keefe et al., 1998; Park et al., 2011) and humans (Suthana et al., 2009; Ekstrom et al., 2003), as well as allocentric view cells in primates (Rolls, 1999, 2023), it would be perhaps be more surprising if we did not see voxels within CA1 that contained scene-sensitive information.

The results from our two analyses are not necessarily at odds, with the subject-level maps demonstrating that scene information is broadly distributed across CA1, which may be a reason for the lack of ‘scene specific’ significant voxels in the group-level analysis. Indeed, meaningful differences between subject-level and group-level analyses are well documented (Jolly & Chang, 2021, Fedorenko, 2021) and can arise due to individual variation in the spatial distrubution of category relevent voxels (Etzel, Zacks, & Braver, 2013). If, as is suggested by our subject-level maps, category information is not represented in as consistent a topological manor in CA1 compared with the subiculum, then this would result in less power to detect an effect at the group level. This individual variation has also been reported in hippocampal MRI connectivity analyses (Borne et al., 2023, Dalton et al., 2022, Przeździk et al., 2019).

Outside of the hippocampus, the face-selective map matched known mapping of face sensitive regions. This was right lateralised (Haxby et al., 2000; Rapcsak, 2019; Rossion, 2014) and also included regions well known to support face processing, such as a portion of the inferior temporal cortex, likely reflecting the fusiform face area, and a portion of the posterior-superior temporal sulcus (Barton, 2022; Kanwisher & Yovel, 2006; Rapcsak, 2019). As mentioned, structural connectivity analysis has shown that these regions connect to the lateral hippocampus (Dalton, 2022) (also reported in our individual level analysis). Whilst our group-level results did not reveal face-selective regions within the hippocampus, this may have been due to high individual variation in the location of face-selective voxels, which we demonstrated were more spatially distributed compared to scene-selective voxels in our subject overlap maps (Figure 2-3). In addition, the nature of our analysis was rather conservative as the construction of the face-selective maps necessarily excluded voxels that were sensitive to more than one category. For example, hippocampal regions that may have been sensitive to both faces and objects would not have been included.

### 4.3 The results align with ideas of parallel processing of category information in networks within, and connected to, the hippocampus

Together considering the differing gradient patterns for scenes and faces from our probabilistic maps (aligning with Dalton et al., 2022 mapping), as well as the scene selectivity patterns in the distal subicular complex and inferior CA1 (aligning with Aggleton, 2012), our interpretations are encompassed by, and build upon, the PMAT (Posterior Medial and an Anterior-Temporal) framework (Inhoff and Ranganath 2017). In this framework, two anatomically distinct parallel processing streams (though see Connor & Knierim, 2017), PM and AT reside in both distributed cortex regions as well as medial temporal lobe regions, the PHC and PrC, respectively, before converging in different regions of the hippocampus, and they support behaviours across a range of cognitive domains (e.g., both mnemonic and perceptual) for different modalities. The PM network is proposed to support episodic simulation, encompassing spatial processing required for our scene oddity task, and the AT network supports representation of item information, encompassing face and object processing required for our face and object oddity tasks. Our results add to this framework by providing insight into the topography of the PM within the hippocampus. We also map aspects of the AT within the cortex, however, as mentioned, our methods to reveal face only information excluded regions which process both face and object information, limiting the ability to accurately map AT within the hippocampus.

### 4.4 Limitations

There are some limitations of our study, and our results should be interpreted with these in mind. First, our small sample size may mean that comparing classifier performance to theoretical chance level (e.g., 50% for 2 class classification) may have led to inflated results, as actual chance level in small sample sizes can exceed theoretical chance level (Combrisson & Jerbi, 2015). Although our conservative group statistics approach reduced this risk, replication of our results with larger sample sizes would be beneficial. Second, we opted to use a high-resolution sequence at 7T to optimize SNR in hippocampal regions, but the FOV only had partial brain coverage. Therefore, we have not mapped all the scene- or face-selective regions that could work in conjunction with the subicular complex and CA1, including regions of entorhinal and perirhinal cortex (Grande et al., 2022; Maass et al., 2015). To map all cortical regions processing category information in conjunction with the subicular complex and CA1, future work with whole-brain high-resolution imaging would be ideal but is challenging with current scanning limitations.

Searchlight methods also have limitations in that they are inherently spatially blurred because the decoding accuracy in each voxel is determined by the information in the surrounding voxels, that is, the ‘neighbourhood’ (Etzel et al., 2013). This means that a significant finding in one voxel and not its neighbour, does not necessarily translate to category information existing in the first voxel but not the second. We therefore do not interpret the results at this voxel-by-voxel level.

Lastly, it is important to note that although we interpret increased scene classification performance in a region as an indication that this region contributes to scene representations, this does not mean that those representations were available to the SVM. For example, it is very unlikely that the scale of scene representations on the neural level matched our sampling of fMRI information. We can conclude that there is decodable information, but we cannot conclude that the patterns providing the decodable information are category representations (Ritchie et al., 2019; Peelen and Downing, 2023). A future step could be to relate MVPA results of a region (e.g., classification performance, distance from classifier decision boundary or pattern similarity) with a behaviour (such as task performance), as such a relationship would provide stronger evidence that the information available to the decoder was a category-specific neural pattern (Ritchie et al., 2019). It was not feasible in this case to relate the searchlight results to behaviour through, for example, correlating with oddity performance, because of the low sample size, high levels of task performance, and low variation in performance across oddity categories.

### 4.5 Conclusions

In conclusion, using MVPA searchlight analyses, in conjunction with high resolution 7T fMRI data from humans performing complex visual concurrent oddity discrimination tasks, our study provides novel support for the importance of the anteromedial (distal) subicular complex but also the inferolateral (proximal) CA1, alongside traditional cortical scene selective regions such as the RSC and PHC, to complex scene discrimination. Therefore, our work has contributed high resolution understanding of hippocampal contributions to the human scene network, demonstrating that analysis across and within hippocampal subfields is crucial for our understanding of hippocampal function and anatomy. In addition, using subject level analysis techniques we demonstrate individual variability of category selective areas within the hippocampus. Finally, contrasting scene selectivity in the medial hippocampus, we provide preliminary evidence for face selectivity along the lateral hippocampus, similar to recent MRI analysis showing lateral hippocampal structural connectivity with face-selective cortical regions, and supporting parallel processing models such as the PMAT framework. Future work could focus on structural and (directed) functional connectivity between these regions to further our understanding of how category information is communicated within category sensitive hippocampal networks.

## Footnotes

1 The medial-lateral axis can also be described at the distal-proximal axis within the subiculum, the regions which border the presubiculum and CA1, respectively. This is also true to an extent in CA1, where medial or distal CA1 borders the subiculum. However, CA1 curves around to border CA2/3 and this is proximal CA1, which is not consistently its most lateral aspect on the anterior-posterior axis.

2 Note that researchers have suggested that this ‘anteromedial subiculum’ scene-selective region detected in fMRI studies may actually correspond to the pre/para-subiculum subregions (Dalton, M. A., & Maguire, E. A. (2017). The pre/parasubiculum: a hippocampal hub for scene-based cognition? *Curr Opin Behav Sci*, *17*, 34-40. https://doi.org/10.1016/j.cobeha.2017.06.001, Ding, S. L. (2013). Comparative anatomy of the prosubiculum, subiculum, presubiculum, postsubiculum, and parasubiculum in human, monkey, and rodent. *J Comp Neurol*, *521*(18), 4145-4162. https://doi.org/10.1002/cne.23416, meaning that the term ‘subicular complex’ may be more appropriate when referring to an ROI that includes multiple subicular subregions (pro-subiculum, subiculum proper, the pre-subiculum, and the para-subiculum). We therefore differentiate between subiculum proper and subiculum complex.

## Acknowledgements

This work was supported by the Biotechnology and Biological Sciences Research Council (BBSRC) [BB/V010549/1; BB/V008242/1; to C.J.H., A.D.L.], the Medical Research Council (G1002149; to K.S.G., C.J.H.), and a Wellcome Trust Strategic Support Fund fellowship (C.J.H.). We would like to thank attendees of British Neuroscience Association Festival of Neuroscience 2023 for their feedback on this work.

## CRediT author statement

Conceptualization: K.S.G., A.D.L., and C.J.H. Data curation: M.-L.R., S.C.B., and C.J.H. Formal analysis: M.-L.R., S.C.B., J.Z., and C.J.H. Funding acquisition: A.D.L. and C.J.H. Investigation: C.J.H. Project administration: M.-L.R., S.C.B., and C.J.H. Supervision: A.D.L. and C.J.H. Visualization: M.-L.R., S.C.B., and C.J.H. Writing - original draft: M.-L.R., S.C.B., and C.J.H. Writing - review & editing: M.-L.R., S.C.B., K.S.G., N.L.V., J.Z., J.P.A., A.D.L., and C.J.H.

## References

Abraham, A., Pedregosa, F., Eickenberg, M., Gervais, P., Mueller, A., Kossaifi, J., … Varoquaux, G. (2014). Machine learning for neuroimaging with scikit-learn. Front Neuroinform, 8, 14. 10.3389/fninf.2014.00014

Aggleton, J. P. (2012). Multiple anatomical systems embedded within the primate medial temporal lobe: implications for hippocampal function. Neurosci Biobehav Rev, 36(7), 1579–1596. 10.1016/j.neubiorev.2011.09.005

Aggleton, J. P., & Christiansen, K. (2015). The subiculum: the heart of the extended hippocampal system. Prog Brain Res, 219, 65–82. 10.1016/bs.pbr.2015.03.003

Aly, M., Ranganath, C., & Yonelinas, A. P. (2013). Detecting changes in scenes: the hippocampus is critical for strength-based perception. Neuron, 78(6), 1127–1137. 10.1016/j.neuron.2013.04.018

Amaral, D. G., Dolorfo, C., & Alvarez-Royo, P. (1991). Organization of CA1 projections to the subiculum: a PHA-L analysis in the rat. Hippocampus, 1(4), 415–435. 10.1002/hipo.450010410

Bakermans, J. J. W., Warren J., Whittington, J. C. R., & Behrens, T. E. J. (2023). Constructing future behaviour in the hippocampal formation through composition and replay. BioRxiv, 2023.04.07.536053. 10.1101/2023.04.07.536053

Baldassano, C., Esteva, A., Fei-Fei, L., & Beck, D. M. (2016). Two Distinct Scene-Processing Networks Connecting Vision and Memory. eNeuro, 3(5). 10.1523/eneuro.0178-16.2016

Barense, M. D., Henson, R. N., Lee, A. C., & Graham, K. S. (2010). Medial temporal lobe activity during complex discrimination of faces, objects, and scenes: Effects of viewpoint. Hippocampus, 20(3), 389–401. 10.1002/hipo.20641

Barton, J. J. S. (2022). Face processing in the temporal lobe. Handb Clin Neurol, 187, 191–210. 10.1016/b978-0-12-823493-8.00019-5

Bernstein, M., & Yovel, G. (2015). Two neural pathways of face processing: A critical evaluation of current models. Neuroscience and biobehavioral reviews, 55, 536–546. 10.1016/j.neubiorev.2015.06.010

Bird, C. M., Vargha-Khadem, F., & Burgess, N. (2008). Impaired memory for scenes but not faces in developmental hippocampal amnesia: a case study. Neuropsychologia, 46(4), 1050–1059. 10.1016/j.neuropsychologia.2007.11.007

Boccara, C. N., Sargolini, F., Thoresen, V. H., Solstad, T., Witter, M. P., Moser, E. I., & Moser, M. B. (2010). Grid cells in pre- and parasubiculum. Nat Neurosci, 13(8), 987–994. 10.1038/nn.2602

Boccia, M., Sulpizio, V., Nemmi, F., Guariglia, C., & Galati, G. (2017). Direct and indirect parieto-medial temporal pathways for spatial navigation in humans: evidence from resting-state functional connectivity. Brain structure & function, 222(4), 1945–1957. 10.1007/s00429-016-1318-6

Bainbridge, W. A., Hall, E. H., & Baker, C. I. (2021). Distinct Representational Structure and Localization for Visual Encoding and Recall during Visual Imagery. Cerebral cortex (New York, N.Y. : 1991), 31(4), 1898–1913. 10.1093/cercor/bhaa329

Behrmann, M., Lee, A. C. H., Geskin, J. Z., Graham, K. S., & Barense, M. D. (2016). Temporal lobe contribution to perceptual function: A tale of three patient groups. Neuropsychologia, 90, 33–45. 10.1016/j.neuropsychologia.2016.05.002

Brun, V. H., Otnass, M. K., Molden, S., Steffenach, H. A., Witter, M. P., Moser, M. B., & Moser, E. I. (2002). Place cells and place recognition maintained by direct entorhinal-hippocampal circuitry. Science (New York, N.Y.), 296(5576), 2243–2246. 10.1126/science.1071089

Brunec, I. K., Bellana, B., Ozubko, J. D., Man, V., Robin, J., Liu, Z. X., … Moscovitch, M. (2018). Multiple Scales of Representation along the Hippocampal Anteroposterior Axis in Humans. Curr Biol, 28(13), 2129–2135.e2126. 10.1016/j.cub.2018.05.016

Bubb, E. J., Kinnavane, L., & Aggleton, J. P. (2017). Hippocampal - diencephalic - cingulate networks for memory and emotion: An anatomical guide. Brain Neurosci Adv, 1(1). 10.1177/2398212817723443

Buckley, M. J., Booth, M. C., Rolls, E. T., & Gaffan, D. (2001). Selective perceptual impairments after perirhinal cortex ablation. J Neurosci, 21(24), 9824–9836.

Chang, C. H., Zehra, S., Nestor, A., & Lee, A. C. (2023). Using image reconstruction to investigate face perception in amnesia. Neuropsychologia, 185, 108573. 10.1016/j.neuropsychologia.2023.108573

Christiansen, K., Metzler-Baddeley, C., Parker, G. D., Muhlert, N., Jones, D. K., Aggleton, J. P., & Vann, S. D. (2017). Topographic separation of fornical fibers associated with the anterior and posterior hippocampus in the human brain: An MRI-diffusion study. Brain Behav, 7(1), e00604. 10.1002/brb3.604

Combrisson, E., & Jerbi, K. (2015). Exceeding chance level by chance: The caveat of theoretical chance levels in brain signal classification and statistical assessment of decoding accuracy. J Neurosci Methods, 250, 126–136. 10.1016/j.jneumeth.2015.01.010

Connor, C. E., & Knierim, J. J. (2017). Integration of objects and space in perception and memory. Nature neuroscience, 20(11), 1493–1503. 10.1038/nn.4657

Cukur, T., Huth, A. G., Nishimoto, S., & Gallant, J. L. (2016). Functional Subdomains within Scene-Selective Cortex: Parahippocampal Place Area, Retrosplenial Complex, and Occipital Place Area. J Neurosci, 36(40), 10257–10273. 10.1523/jneurosci.4033-14.2016

Dalton, M. A., D’Souza, A., Lv, J., & Calamante, F. (2022). New insights into anatomical connectivity along the anterior-posterior axis of the human hippocampus using in vivo quantitative fibre tracking. Elife, 11. 10.7554/eLife.76143

Dalton, M. A., & Maguire, E. A. (2017). The pre/parasubiculum: a hippocampal hub for scene-based cognition? Curr Opin Behav Sci, 17, 34–40. 10.1016/j.cobeha.2017.06.001

Dalton, M. A., McCormick, C., & Maguire, E. A. (2019). Differences in functional connectivity along the anterior-posterior axis of human hippocampal subfields. Neuroimage, 192, 38–51. 10.1016/j.neuroimage.2019.02.066

Ding, S. L. (2013). Comparative anatomy of the prosubiculum, subiculum, presubiculum, postsubiculum, and parasubiculum in human, monkey, and rodent. J Comp Neurol, 521(18), 4145–4162. 10.1002/cne.23416

Ekstrom, A. D., Kahana, M. J., Caplan, J. B., Fields, T. A., Isham, E. A., Newman, E. L., & Fried, I. (2003). Cellular networks underlying human spatial navigation. Nature, 425(6954), 184–188. 10.1038/nature01964

Epstein, R. A. (2008). Parahippocampal and retrosplenial contributions to human spatial navigation. Trends Cogn Sci, 12(10), 388–396. 10.1016/j.tics.2008.07.004

Epstein, R. A., & Baker, C. I. (2019). Scene Perception in the Human Brain. Annu Rev Vis Sci, 5, 373–397. 10.1146/annurev-vision-091718-014809

Epstein, R. A., Patai, E. Z., Julian, J. B., & Spiers, H. J. (2017). The cognitive map in humans: spatial navigation and beyond. Nat Neurosci, 20(11), 1504–1513. 10.1038/nn.4656

Erez, J., Lee, A. C., & Barense, M. D. (2013). It does not look odd to me: perceptual impairments and eye movements in amnesic patients with medial temporal lobe damage. Neuropsychologia, 51(1), 168–180. 10.1016/j.neuropsychologia.2012.11.003

Etzel, J. A., Zacks, J. M., & Braver, T. S. (2013). Searchlight analysis: promise, pitfalls, and potential. Neuroimage, 78, 261–269. 10.1016/j.neuroimage.2013.03.041

Evensmoen, H. R., Lehn, H., Xu, J., Witter, M. P., Nadel, L., & Håberg, A. K. (2013). The anterior hippocampus supports a coarse, global environmental representation and the posterior hippocampus supports fine-grained, local environmental representations. J Cogn Neurosci, 25(11), 1908–1925. 10.1162/jocn_a_00436

Ezama, L., Hernández-Cabrera, J. A., Seoane, S., Pereda, E., & Janssen, N. (2021). Functional connectivity of the hippocampus and its subfields in resting-state networks. Eur J Neurosci, 53(10), 3378–3393. 10.1111/ejn.15213

Fedorenko, E. (2021). The early origins and the growing popularity of the individual-subject analytic approach in human neuroscience. Deep Imaging - Personalized Neuroscience, 40, 105–112. 10.1016/j.cobeha.2021.02.023

Fritch, H. A., Spets, D. S., & Slotnick, S. D. (2021). Functional connectivity with the anterior and posterior hippocampus during spatial memory. Hippocampus, 31(7), 658–668. 10.1002/hipo.23283

Frost, B. E., Martin, S. K., Cafalchio, M., Islam, M. N., Aggleton, J. P., & O’Mara, S. M. (2021). Anterior Thalamic Inputs Are Required for Subiculum Spatial Coding, with Associated Consequences for Hippocampal Spatial Memory. J Neurosci, 41(30), 6511–6525. 10.1523/jneurosci.2868-20.2021

Gaffan, D. (1992). Amnesia for Complex Naturalistic Scenes and for Objects Following Fornix Transection in the Rhesus Monkey. Eur J Neurosci, 4(5), 381–388. 10.1111/j.1460-9568.1992.tb00886.x

Gardette, J., Cousin, E., Bourgin, J., Torlay, L., Pichat, C., Moreaud, O., & Hot, P. (2022). Hippocampal activity during memory and visual perception: The role of representational content. Cortex; a journal devoted to the study of the nervous system and behavior, 157, 14–29. 10.1016/j.cortex.2022.09.004

Gardette, J., Mosca, C., Asien, C., Borg, C., Mazzola, L., Convers, P., … & Hot, P. (2023). Complex visual discrimination is impaired after right, but not left, anterior temporal lobectomy. Hippocampus.

Genon, S., Bernhardt, B. C., La Joie, R., Amunts, K., & Eickhoff, S. B. (2021). The many dimensions of human hippocampal organization and (dys)function. Trends Neurosci, 44(12), 977–989. 10.1016/j.tins.2021.10.003

Gigg, J., Finch, D. M., & O’Mara, S. M. (2000). Responses of rat subicular neurons to convergent stimulation of lateral entorhinal cortex and CA1 in vivo. Brain Res, 884(1--2), 35–50. 10.1016/s0006-8993(00)02878-x

Gigg, J. (2006). Constraints on hippocampal processing imposed by the connectivity between CA1, subiculum and subicular targets. Behavioural brain research, 174(2), 265–271. 10.1016/j.bbr.2006.06.014

Goodale, M. A., & Milner, A. D. (1992). Separate visual pathways for perception and action. Trends in neurosciences, 15(1), 20–25. 10.1016/0166-2236(92)90344-8

Grill-Spector, K., Weiner, K. S., Kay, K., & Gomez, J. (2017). The Functional Neuroanatomy of Human Face Perception. Annual review of vision science, 3, 167–196. 10.1146/annurev-vision-102016-061214

Grande, X., Sauvage, M. M., Becke, A., Düzel, E., & Berron, D. (2022). Transversal functional connectivity and scene-specific processing in the human entorhinal-hippocampal circuitry. Elife, 11. 10.7554/eLife.76479

Guest, O., & Love, B. C. (2017). What the success of brain imaging implies about the neural code. Elife, 6. 10.7554/eLife.21397

Hartley, T., Bird, C. M., Chan, D., Cipolotti, L., Husain, M., Vargha-Khadem, F., & Burgess, N. (2007). The hippocampus is required for short-term topographical memory in humans. Hippocampus, 17(1), 34–48. 10.1002/hipo.20240

Haxby, J. V., Connolly, A. C., & Guntupalli, J. S. (2014). Decoding neural representational spaces using multivariate pattern analysis. Annual review of neuroscience, 37, 435–456. 10.1146/annurev-neuro-062012-170325

Haxby J. V. (2012). Multivariate pattern analysis of fMRI: the early beginnings. NeuroImage, 62(2), 852–855. 10.1016/j.neuroimage.2012.03.016

Haxby, J. V., Hoffman, E. A., & Gobbini, M. I. (2000). The distributed human neural system for face perception. Trends Cogn Sci, 4(6), 223–233.

Henriksen, E. J., Colgin, L. L., Barnes, C. A., Witter, M. P., Moser, M. B., & Moser, E. I. (2010). Spatial representation along the proximodistal axis of CA1. Neuron, 68(1), 127–137. 10.1016/j.neuron.2010.08.042

Hodgetts, C. J., Postans, M., Shine, J. P., Jones, D. K., Lawrence, A. D., & Graham, K. S. (2015). Dissociable roles of the inferior longitudinal fasciculus and fornix in face and place perception. Elife, 4. 10.7554/eLife.07902

Hodgetts, C. J., Shine, J. P., Lawrence, A. D., Downing, P. E., & Graham, K. S. (2016). Evidencing a place for the hippocampus within the core scene processing network. Human Brain Mapping, 37(11), 3779–3794. 10.1002/hbm.23275

Hodgetts, C. J., Voets, N. L., Thomas, A. G., Clare, S., Lawrence, A. D., & Graham, K. S. (2017). Ultra-High-Field fMRI Reveals a Role for the Subiculum in Scene Perceptual Discrimination. J Neurosci, 37(12), 3150–3159. 10.1523/jneurosci.3225-16.2017

Honda, Y., & Shibata, H. (2017). Organizational connectivity among the CA1, subiculum, presubiculum, and entorhinal cortex in the rabbit. J Comp Neurol, 525(17), 3705–3741. 10.1002/cne.24297

Honda, Y., Shimokawa, T., Matsuda, S., Kobayashi, Y., & Moriya-Ito, K. (2022). Hippocampal Connectivity of the Presubiculum in the Common Marmoset (Callithrix jacchus). Front Neural Circuits, 16, 863478. 10.3389/fncir.2022.863478

Huang, C. C., Rolls, E. T., Hsu, C. H., Feng, J., & Lin, C. P. (2021). Extensive Cortical Connectivity of the Human Hippocampal Memory System: Beyond the “What” and “Where” Dual Stream Model. Cerebral cortex (New York, N.Y. : 1991), 31(10), 4652–4669. 10.1093/cercor/bhab113

Jenkinson, M., Bannister, P., Brady, M., & Smith, S. (2002). Improved optimization for the robust and accurate linear registration and motion correction of brain images. Neuroimage, 17(2), 825–841. 10.1016/s1053-8119(02)91132-8

Jenkinson, M., Beckmann, C. F., Behrens, T. E., Woolrich, M. W., & Smith, S. M. (2012). FSL. Neuroimage, 62(2), 782–790. 10.1016/j.neuroimage.2011.09.015

Jolly, E., & Chang, L. J. (2021). Multivariate spatial feature selection in fMRI. Social cognitive and affective neuroscience, 16(8), 795–806. 10.1093/scan/nsab010

Kanwisher, N., & Yovel, G. (2006). The fusiform face area: a cortical region specialized for the perception of faces. Philos Trans R Soc Lond B Biol Sci, 361(1476), 2109–2128. 10.1098/rstb.2006.1934

Kim, S. M., Ganguli, S., & Frank, L. M. (2012). Spatial information outflow from the hippocampal circuit: distributed spatial coding and phase precession in the subiculum. J Neurosci, 32(34), 11539–11558. 10.1523/jneurosci.5942-11.2012

Kloosterman, F., Van Haeften, T., Witter, M. P., & Lopes Da Silva, F. H. (2003). Electrophysiological characterization of interlaminar entorhinal connections: an essential link for re-entrance in the hippocampal-entorhinal system. The European journal of neuroscience, 18(11), 3037–3052. 10.1111/j.1460-9568.2003.03046.x

Kravitz, D. J., Saleem, K. S., Baker, C. I., & Mishkin, M. (2011). A new neural framework for visuospatial processing. Nat Rev Neurosci, 12(4), 217–230. 10.1038/nrn3008

Kriegeskorte, N., Mur, M., Ruff, D. A., Kiani, R., Bodurka, J., Esteky, H., … Bandettini, P. A. (2008). Matching categorical object representations in inferior temporal cortex of man and monkey. Neuron, 60(6), 1126–1141. 10.1016/j.neuron.2008.10.043

Ledergerber, D., Battistin, C., Blackstad, J. S., Gardner, R. J., Witter, M. P., Moser, M. B., … Moser, E. I. (2021). Task-dependent mixed selectivity in the subiculum. Cell Rep, 35(8), 109175. 10.1016/j.celrep.2021.109175

Lee, A. C., Brodersen, K. H., & Rudebeck, S. R. (2013). Disentangling spatial perception and spatial memory in the hippocampus: a univariate and multivariate pattern analysis fMRI study. J Cogn Neurosci, 25(4), 534–546. 10.1162/jocn_a_00301

Lee, A. C., Buckley, M. J., Gaffan, D., Emery, T., Hodges, J. R., & Graham, K. S. (2006). Differentiating the roles of the hippocampus and perirhinal cortex in processes beyond long-term declarative memory: a double dissociation in dementia. J Neurosci, 26(19), 5198–5203. 10.1523/jneurosci.3157-05.2006

Lee, A. C., Buckley, M. J., Pegman, S. J., Spiers, H., Scahill, V. L., Gaffan, D., … Graham, K. S. (2005). Specialization in the medial temporal lobe for processing of objects and scenes. Hippocampus, 15(6), 782–797. 10.1002/hipo.20101

Lee, A. C., Scahill, V. L., & Graham, K. S. (2008). Activating the medial temporal lobe during oddity judgment for faces and scenes. Cereb Cortex, 18(3), 683–696. 10.1093/cercor/bhm104

Lee, A. C., Yeung, L. K., & Barense, M. D. (2012). The hippocampus and visual perception. Front Hum Neurosci, 6, 91. 10.3389/fnhum.2012.00091

Leutgeb, J. K., Leutgeb, S., Moser, M. B., & Moser, E. I. (2007). Pattern separation in the dentate gyrus and CA3 of the hippocampus. Science, 315(5814), 961–966. 10.1126/science.1135801

Lever, C., Burton, S., Jeewajee, A., O’Keefe, J., & Burgess, N. (2009). Boundary vector cells in the subiculum of the hippocampal formation. J Neurosci, 29(31), 9771–9777. 10.1523/jneurosci.1319-09.2009

Liang, J. C., Wagner, A. D., & Preston, A. R. (2013). Content representation in the human medial temporal lobe. Cereb Cortex, 23(1), 80–96. 10.1093/cercor/bhr379

Maass, A., Berron, D., Libby, L. A., Ranganath, C., & Düzel, E. (2015). Functional subregions of the human entorhinal cortex. Elife, 4. 10.7554/eLife.06426

Margulies, D. S., Vincent, J. L., Kelly, C., Lohmann, G., Uddin, L. Q., Biswal, B. B., Villringer, A., Castellanos, F. X., Milham, M. P., & Petrides, M. (2009). Precuneus shares intrinsic functional architecture in humans and monkeys. Proceedings of the National Academy of Sciences of the United States of America, 106(47), 20069–20074. 10.1073/pnas.0905314106

Maguire, E. A., & Mullally, S. L. (2013). The hippocampus: a manifesto for change. J Exp Psychol Gen, 142(4), 1180–1189. 10.1037/a0033650

Martin, C. B., & Barense, M. D. (2023). Perception and Memory in the Ventral Visual Stream and Medial Temporal Lobe. Annual review of vision science, 9, 409–434. 10.1146/annurev-vision-120222-014200

Matsumoto, N., Kitanishi, T., & Mizuseki, K. (2019). The subiculum: Unique hippocampal hub and more. Neurosci Res, 143, 1–12. 10.1016/j.neures.2018.08.002

Mayes, A., Montaldi, D., & Migo, E. (2007). Associative memory and the medial temporal lobes. Trends in cognitive sciences, 11(3), 126–135. 10.1016/j.tics.2006.12.003

McCormick, C., Dalton, M. A., Zeidman, P., & Maguire, E. A. (2021). Characterising the hippocampal response to perception, construction and complexity. Cortex; a journal devoted to the study of the nervous system and behavior, 137, 1–17. 10.1016/j.cortex.2020.12.018

McGugin, R. W., Van Gulick, A. E., & Gauthier, I. (2016). Cortical Thickness in Fusiform Face Area Predicts Face and Object Recognition Performance. J Cogn Neurosci, 28(2), 282–294. 10.1162/jocn_a_00891

Mizuseki, K., & Kitanishi, T. (2022). Oscillation-coordinated, noise-resistant information distribution via the subiculum. Curr Opin Neurobiol, 75, 102556. 10.1016/j.conb.2022.102556

Murray, E. A., Wise, S. P., & Graham, K. S. (2017). The Evolution of Memory Systems: Ancestors, Anatomy, and Adaptations. Oxford University Press.

Murray, E. A., Wise, S. P., & Graham, K. S. (2018). Representational specializations of the hippocampus in phylogenetic perspective. Neurosci Lett, 680, 4–12. 10.1016/j.neulet.2017.04.065

Naber, P. A., & Witter, M. P. (1998). Subicular efferents are organized mostly as parallel projections: a double-labeling, retrograde-tracing study in the rat. J Comp Neurol, 393(3), 284–297.

Nadel, L., Hoscheidt, S., & Ryan, L. R. (2013). Spatial cognition and the hippocampus: the anterior-posterior axis. J Cogn Neurosci, 25(1), 22–28. 10.1162/jocn_a_00313

Nasr, S., Devaney, K. J., & Tootell, R. B. (2013). Spatial encoding and underlying circuitry in scene-selective cortex. Neuroimage, 83, 892–900. 10.1016/j.neuroimage.2013.07.030

Navarro Schröder, T., Haak, K. V., Zaragoza Jimenez, N. I., Beckmann, C. F., & Doeller, C. F. (2015). Functional topography of the human entorhinal cortex. Elife, 4. 10.7554/eLife.06738

Ng, C. W., Elias, G. A., Asem, J. S. A., Allen, T. A., & Fortin, N. J. (2018). Nonspatial sequence coding varies along the CA1 transverse axis. Behav Brain Res, 354, 39–47. 10.1016/j.bbr.2017.10.015

Norman, K. A., Polyn, S. M., Detre, G. J., & Haxby, J. V. (2006). Beyond mind-reading: multi-voxel pattern analysis of fMRI data. Trends Cogn Sci, 10(9), 424–430. 10.1016/j.tics.2006.07.005

O’Keefe, J., Burgess, N., Donnett, J. G., Jeffery, K. J., & Maguire, E. A. (1998). Place cells, navigational accuracy, and the human hippocampus. Philosophical transactions of the Royal Society of London. Series B, Biological sciences, 353(1373), 1333–1340. 10.1098/rstb.1998.0287

O’Keefe, J., & Recce, M. L. (1993). Phase relationship between hippocampal place units and the EEG theta rhythm. Hippocampus, 3(3), 317–330. 10.1002/hipo.450030307

O’Mara, S. (2005). The subiculum: what it does, what it might do, and what neuroanatomy has yet to tell us. J Anat, 207(3), 271–282. 10.1111/j.1469-7580.2005.00446.x

O’Mara, S. M., & Aggleton, J. P. (2019). Space and Memory (Far) Beyond the Hippocampus: Many Subcortical Structures Also Support Cognitive Mapping and Mnemonic Processing. Front Neural Circuits, 13, 52. 10.3389/fncir.2019.00052

O’Mara, S. M., Sanchez-Vives, M. V., Brotons-Mas, J. R., & O’Hare, E. (2009). Roles for the subiculum in spatial information processing, memory, motivation and the temporal control of behaviour. Prog Neuropsychopharmacol Biol Psychiatry, 33(5), 782–790. 10.1016/j.pnpbp.2009.03.040

Oliva, A., Fernández-Ruiz, A., Buzsáki, G., & Berényi, A. (2016). Spatial coding and physiological properties of hippocampal neurons in the Cornu Ammonis subregions. Hippocampus, 26(12), 1593–1607. 10.1002/hipo.22659

Olson, J. M., Tongprasearth, K., & Nitz, D. A. (2017). Subiculum neurons map the current axis of travel. Nat Neurosci, 20(2), 170–172. 10.1038/nn.4464

Olsen, R. K., Moses, S. N., Riggs, L., & Ryan, J. D. (2012). The hippocampus supports multiple cognitive processes through relational binding and comparison. Frontiers in human neuroscience, 6, 146. 10.3389/fnhum.2012.00146

Park, E., Dvorak, D., & Fenton, A. A. (2011). Ensemble place codes in hippocampus: CA1, CA3, and dentate gyrus place cells have multiple place fields in large environments. PloS one, 6(7), e22349. 10.1371/journal.pone.0022349

Peelen, M. V., & Downing, P. E. (2023). Testing cognitive theories with multivariate pattern analysis of neuroimaging data. Nature human behaviour, 7(9), 1430–1441. 10.1038/s41562-023-01680-z

Preston, A. R., Bornstein, A. M., Hutchinson, J. B., Gaare, M. E., Glover, G. H., & Wagner, A. D. (2010). High-resolution fMRI of content-sensitive subsequent memory responses in human medial temporal lobe. J Cogn Neurosci, 22(1), 156–173. 10.1162/jocn.2009.21195

Preston-Ferrer, P., Coletta, S., Frey, M., & Burgalossi, A. (2016). Anatomical organization of presubicular head-direction circuits. Elife, 5. 10.7554/eLife.14592

Rapcsak, S. Z. (2019). Face Recognition. Curr Neurol Neurosci Rep, 19(7), 41. 10.1007/s11910-019-0960-9

Reid, A. T., Headley, D. B., Mill, R. D., Sanchez-Romero, R., Uddin, L. Q., Marinazzo, D., Lurie, D. J., Valdés-Sosa, P. A., Hanson, S. J., Biswal, B. B., Calhoun, V., Poldrack, R. A., & Cole, M. W. (2019). Advancing functional connectivity research from association to causation. Nature neuroscience, 22(11), 1751–1760. 10.1038/s41593-019-0510-4

Ritchie, J. B., Kaplan, D. M., & Klein, C. (2019). Decoding the Brain: Neural Representation and the Limits of Multivariate Pattern Analysis in Cognitive Neuroscience. Br J Philos Sci, 70(2), 581–607. 10.1093/bjps/axx023

Robertson, R. G., Rolls, E. T., & Georges-François, P. (1998). Spatial view cells in the primate hippocampus: effects of removal of view details. Journal of neurophysiology, 79(3), 1145–1156. 10.1152/jn.1998.79.3.1145

Robertson, R. G., Rolls, E. T., Georges-François, P., & Panzeri, S. (1999). Head direction cells in the primate pre-subiculum. Hippocampus, 9(3), 206–219. 10.1002/(sici)1098-1063(1999)9:3<206::aid-hipo2>3.0.co;2-h

Rolls E. T. (1999). Spatial view cells and the representation of place in the primate hippocampus. Hippocampus, 9(4), 467–480. 10.1002/(SICI)1098-1063(1999)9:4<467::AID-HIPO13>3.0.CO;2-F

Rolls, E. T. (2021). Neurons including hippocampal spatial view cells, and navigation in primates including humans. Hippocampus, 31(6), 593–611. 10.1002/hipo.23324

Rolls, E. T. (2023). Hippocampal spatial view cells for memory and navigation, and their underlying connectivity in humans. Hippocampus, 33(5), 533–572. 10.1002/hipo.23467

Rossion, B. (2014). Understanding face perception by means of prosopagnosia and neuroimaging. Front Biosci (Elite Ed), 6(2), 258–307. 10.2741/e706

Schultz, C., & Engelhardt, M. (2014). Anatomy of the hippocampal formation. Front Neurol Neurosci, 34, 6–17. 10.1159/000360925

Schultz, H., Sommer, T., & Peters, J. (2015). The Role of the Human Entorhinal Cortex in a Representational Account of Memory. Frontiers in human neuroscience, 9, 628. 10.3389/fnhum.2015.00628

Sekeres, M. J., Winocur, G., & Moscovitch, M. (2018). The hippocampus and related neocortical structures in memory transformation. Neurosci Lett, 680, 39–53. 10.1016/j.neulet.2018.05.006

Sharma, A., Nair, I. R., & Yoganarasimha, D. (2022). Attractor-like Dynamics in the Subicular Complex. The Journal of neuroscience : the official journal of the Society for Neuroscience, 42(40), 7594–7614. 10.1523/JNEUROSCI.2048-20.2022

Sharp, P. E. (2006). Subicular place cells generate the same “map” for different environments: comparison with hippocampal cells. Behav Brain Res, 174(2), 206–214. 10.1016/j.bbr.2006.05.034

Shine, J. P., Hodgetts, C. J., Postans, M., Lawrence, A. D., & Graham, K. S. (2015). APOE-epsilon4 selectively modulates posteromedial cortex activity during scene perception and short-term memory in young healthy adults. Sci Rep, 5, 16322. 10.1038/srep16322

Simonsen, Ø. W., Czajkowski, R., & Witter, M. P. (2022). Retrosplenial and subicular inputs converge on superficially projecting layer V neurons of medial entorhinal cortex. Brain structure & function, 227(8), 2821–2837. 10.1007/s00429-022-02578-8

Smith, S. M. (2002). Fast robust automated brain extraction. Hum Brain Mapp, 17(3), 143–155. 10.1002/hbm.10062

Smith, S. M., & Nichols, T. E. (2009). Threshold-free cluster enhancement: addressing problems of smoothing, threshold dependence and localisation in cluster inference. NeuroImage, 44(1), 83–98. 10.1016/j.neuroimage.2008.03.061

Soto, D., Greene, C. M., Kiyonaga, A., Rosenthal, C. R., & Egner, T. (2012). A parieto-medial temporal pathway for the strategic control over working memory biases in human visual attention. The Journal of neuroscience : the official journal of the Society for Neuroscience, 32(49), 17563–17571. 10.1523/JNEUROSCI.2647-12.2012

Stefanini, F., Kushnir, L., Jimenez, J. C., Jennings, J. H., Woods, N. I., Stuber, G. D., … Fusi, S. (2020). A Distributed Neural Code in the Dentate Gyrus and in CA1. Neuron, 107(4), 703–716.e704. 10.1016/j.neuron.2020.05.022

Strange, B. A., Witter, M. P., Lein, E. S., & Moser, E. I. (2014). Functional organization of the hippocampal longitudinal axis. Nat Rev Neurosci, 15(10), 655–669. 10.1038/nrn3785

Sun, Y., Nitz, D. A., Xu, X., & Giocomo, L. M. (2023). The subiculum encodes environmental geometry. BioRxiv, 2023.05.07.539721. 10.1101/2023.05.07.539721

Suthana, N. A., Ekstrom, A. D., Moshirvaziri, S., Knowlton, B., & Bookheimer, S. Y. (2009). Human hippocampal CA1 involvement during allocentric encoding of spatial information. The Journal of neuroscience : the official journal of the Society for Neuroscience, 29(34), 10512–10519. 10.1523/JNEUROSCI.0621-09.2009

Taylor, K. J., Henson, R. N., & Graham, K. S. (2007). Recognition memory for faces and scenes in amnesia: dissociable roles of medial temporal lobe structures. Neuropsychologia, 45(11), 2428–2438. 10.1016/j.neuropsychologia.2007.04.004

Thompson, P. M., Jahanshad, N., Ching, C. R. K., Salminen, L. E., Thomopoulos, S. I., Bright, J., … Zelman, V. (2020). ENIGMA and global neuroscience: A decade of large-scale studies of the brain in health and disease across more than 40 countries. Transl Psychiatry, 10(1), 100. 10.1038/s41398-020-0705-1

Treder, M. S. (2020). MVPA-Light: A Classification and Regression Toolbox for Multi-Dimensional Data. Front Neurosci, 14, 289. 10.3389/fnins.2020.00289

Turk-Browne, N. B. (2019). The hippocampus as a visual area organized by space and time: A spatiotemporal similarity hypothesis. Vision Res, 165, 123–130. 10.1016/j.visres.2019.10.007

Vogel, J. W., La Joie, R., Grothe, M. J., Diaz-Papkovich, A., Doyle, A., Vachon-Presseau, E., Lepage, C., Vos de Wael, R., Thomas, R. A., Iturria-Medina, Y., Bernhardt, B., Rabinovici, G. D., & Evans, A. C. (2020). A molecular gradient along the longitudinal axis of the human hippocampus informs large-scale behavioral systems. Nature communications, 11(1), 960. 10.1038/s41467-020-14518-3

Weaverdyck, M. E., Lieberman, M. D., & Parkinson, C. (2020). Tools of the Trade Multivoxel pattern analysis in fMRI: a practical introduction for social and affective neuroscientists. Soc Cogn Affect Neurosci, 15(4), 487–509. 10.1093/scan/nsaa057

Winkler, A. M., Ridgway, G. R., Webster, M. A., Smith, S. M., & Nichols, T. E. (2014). Permutation inference for the general linear model. Neuroimage, 92(100), 381–397. 10.1016/j.neuroimage.2014.01.060

Wirt, R. A., & Hyman, J. M. (2017). Integrating Spatial Working Memory and Remote Memory: Interactions between the Medial Prefrontal Cortex and Hippocampus. Brain Sci, 7(4). 10.3390/brainsci7040043

Witter M. P. (2006). Connections of the subiculum of the rat: topography in relation to columnar and laminar organization. Behavioural brain research, 174(2), 251–264. 10.1016/j.bbr.2006.06.022

Witter, M. P., & Amaral, D. G. (2021). The entorhinal cortex of the monkey: VI. Organization of projections from the hippocampus, subiculum, presubiculum, and parasubiculum. The Journal of comparative neurology, 529(4), 828–852. 10.1002/cne.24983

Woolrich, M. W., Ripley, B. D., Brady, M., & Smith, S. M. (2001). Temporal autocorrelation in univariate linear modeling of FMRI data. Neuroimage, 14(6), 1370–1386. 10.1006/nimg.2001.0931

Yang, Z., Fang, F., & Weng, X. (2012). Recent developments in multivariate pattern analysis for functional MRI. Neurosci Bull, 28(4), 399–408. 10.1007/s12264-012-1253-3

Zeidman, P., Lutti, A., & Maguire, E. A. (2015). Investigating the functions of subregions within anterior hippocampus. Cortex, 73, 240–256. 10.1016/j.cortex.2015.09.002

Zeidman, P., & Maguire, E. A. (2016). Anterior hippocampus: the anatomy of perception, imagination and episodic memory. In Nat Rev Neurosci (Vol. 17, pp. 173–182). 10.1038/nrn.2015.24

Zeidman, P., Mullally, S. L., & Maguire, E. A. (2015). Constructing, Perceiving, and Maintaining Scenes: Hippocampal Activity and Connectivity. Cereb Cortex, 25(10), 3836–3855. 10.1093/cercor/bhu266

Ziegler, M. G., Liu, Z. X., Arsenault, J., Dang, C., Grady, C., Rosenbaum, R. S., & Moscovitch, M. (2023). Differential involvement of the anterior and posterior hippocampus, parahippocampus, and retrosplenial cortex in making precise judgments of spatial distance and object size for remotely acquired memories of environments and objects. Cerebral cortex (New York, N.Y. : 1991), 33(18), 10139–10154. 10.1093/cercor/bhad272

